# Novel tool use does not depend on mechanical reasoning: evidence from apraxia

**DOI:** 10.64898/2026.05.14.724638

**Authors:** Yue Du, Simon Thibault, John Yates, Laurel J. Buxbaum, John W. Krakauer, Aaron L. Wong

## Abstract

A hallmark of human intelligence is the ability to use tools. Yet the cognitive processes supporting this ability remain debated. One contemporary view holds that mechanical reasoning is central for tool use, especially in the case of tools with which we have no prior experience. However, previous support for the role of mechanical reasoning often relies on circular logic, wherein poor performance on novel tool-use tasks is taken as evidence that impaired mechanical reasoning causes tool-use deficits in limb apraxia. To address this limitation, we independently assessed mechanical reasoning and novel tool use in separate tasks in individuals with limb apraxia, and compared their performance to individuals without apraxia. We also examined whether these two abilities are similarly associated with other cognitive abilities including motor imagery, mental rotation of non-body objects, general reasoning, and spatial working memory. Finally, we explored brain-behavior relationships using support vector regression lesion-symptom mapping. Our behavioral and imaging data together showed that mechanical reasoning does not underlie novel tool-use deficits in apraxia. Graphical analysis further revealed that novel tool use and mechanical reasoning loaded onto distinct latent clusters: novel tool use was strongly associated with other praxis abilities yet separable from cognitive abilities that require reasoning and mental simulation, whereas mechanical reasoning was primarily linked to other high-level reasoning abilities but not tool use. These findings challenge the notion that mechanical reasoning is central to tool-use ability, and instead suggest that tool use is more likely to be an intuitive or automatic process.

## Introduction

Tool use is fundamental to human life. Whether it is something like a hammer for simple housework or advanced technologies like surgical robotics for professional jobs, tools enable us to interact with the world in adaptive and flexible ways (Derex et al., 2019; Mangalam et al., 2022; Osiurak, Lasserre, et al., 2021; Osiurak & Reynaud, 2020). As technology continues to rapidly advance, understanding how people learn and use new tools becomes increasingly critical. Despite a large body of prior research (Goldenberg & Spatt, 2009; Johnson-Frey, 2004; Mangalam et al., 2022; Orban & Caruana, 2014; Osiurak et al., 2013; Vaesen, 2012), the cognitive processes that support using novel tools remain poorly understood.

One account posits that novel tool use relies on mechanical reasoning (Baumard et al., 2018; Osiurak & Badets, 2016) ‒ deliberate mental simulation of how one mechanical component (e.g., the tool) causally affects another (e.g., the recipient) (Battaglia et al., 2013; De Kleer & Brown, 1984; Goldenberg & Hagmann, 1998; Hegarty, 2004; Schwartz & Black, 1996a; Ullman et al., 2017; Weinberg, 2019). Mechanical reasoning is thought to be especially critical for the step of selecting the appropriate tool to achieve a desired outcome. A classic test that is commonly thought to involve mechanical reasoning, for example, asks participants to select and apply the correct hook to lift a cylindrical object (Goldenberg & Hagmann, 1998). In particular, tool selection may require consideration of the shape complementarity between the tool and the cylinder (e.g., does this hook fit into that hole) and their physical interactions (e.g., will the coupling between hook and hole be stable enough to lift the object off the table). These tool-object interactions, in turn, are posited to constrain the body-tool relationship, that is, the proper motor actions to use the tool (Baumard et al., 2021). Thus, mechanical reasoning has been posited as essential for selecting the appropriate tool to perform an action, especially when encountering novel objects with which we have no prior experience (Buxbaum, 2017; Goldenberg & Hagmann, 1998; Osiurak et al., 2009) or when repurposing familiar tools for new uses (Allen et al., 2020), although some have argued that it is central to all tool-based interactions regardless of familiarity (Baumard et al., 2018, 2021; Mangalam et al., 2022). Regardless, this view of mechanical reasoning aligns with the broader idea that people can infer the outcomes of dynamic physical interactions such as the consequences of collisions between objects (Allen et al., 2020; Hamrick et al., 2016; Ullman et al., 2017), and that there is a high degree of overlap between the brain networks involved in physical inference and tool use (Fischer et al., 2016; Fischer & Mahon, 2021).

Although the idea that mechanical reasoning supports novel tool use is intuitively appealing, much of the existing support for this account unfortunately suffers from circular logic. That is, evidence that novel tool use, especially tool selection, depends on mechanical reasoning is largely based on performance in single tasks that are assumed to measure both abilities. In particular, a large body of work examines the performance of individuals with limb apraxia, a cognitive-motor disorder impairing the ability to use tools that arises in about 50% of individuals who have a stroke in the left hemisphere (Goldenberg, 2013b). Apraxia provides a compelling model for investigating which underlying cognitive processes fail when tool use is impaired, as people with apraxia often struggle to select and apply novel tools to perform tasks like retrieving an object from a container (Baumard et al., 2014; Goldenberg & Hagmann, 1998; Jarry et al., 2013; Lesourd et al., 2019; Spatt et al., 2002; Stoll et al., 2022). These deficits in figuring out how to solve the task – particularly in selecting the proper tool to achieve the task goal – have been taken as evidence of impaired mechanical reasoning in these individuals. Yet the same findings have simultaneously been used to claim that impaired mechanical reasoning causes novel tool-use deficits (Baumard et al., 2014, 2016, 2021; Faye et al., 2018, 2018; Goldenberg & Hagmann, 1998; Hodges et al., 1999; Jarry et al., 2013; Osiurak et al., 2009; Spatt et al., 2002). This inversion of the explanandum and explanans adds no independent support for the mechanical reasoning account and just reiterates the original assumption that mechanical reasoning is essential in tool use.

To address this limitation, we measured mechanical reasoning and novel tool use in separate tasks, allowing us to empirically test, rather than assume, the relationship between them. This study was developed to examine two questions. We first assessed whether mechanical reasoning ability is impaired in limb apraxia and is associated with performance of a well-validated novel tool-use task (Goldenberg & Hagmann, 1998). To address this, we developed a mechanical reasoning task inspired by Rube Goldberg machines: multi-step systems comprising simple devices that together perform basic tasks in an overly complicated fashion. In this same spirit, our task required participants to reason about a mechanical system composed of interacting ramps, gears, levers, and/or pulleys – components that form the basis of many real-world tools (Hegarty, 1992, 2004; Hegarty et al., 1988, 2005; Rozenblit et al., 2002; Schwartz & Black, 1996a, 1996b; Yoon & Narayanan, 2004). Importantly, the task is entirely agent-free, requiring no tool selection, motor execution of tool-use actions, or consideration of human-machine interaction. If mechanical reasoning truly underlies tool-use impairments, participants with limb apraxia should perform poorly in this task. Moreover, if the novel tool-use task employed in prior studies indeed depends on mechanical reasoning, we would expect performance on these two tasks to tightly correlate.

Second, we addressed the question of whether mechanical reasoning and novel tool use, especially tool selection, are associated with the same or distinct cognitive processes. That is, if the two are related, they should be show a similar pattern of associations with other cognitive capacities. In contrast, distinct patterns of associations would suggest functional dissociation. To test this possibility, in the same group of participants we assessed several cognitive processes that have previously been thought to be related to mechanical reasoning, tool-use, and/or praxis. One such process is motor imagery, which we tested using a task that has been proposed to require the mental simulation (but not execution) of one’s own hand movements (Parsons, 1994). Motor imagery has previously been linked to mechanical reasoning due to a presumed shared reliance on mental simulation (Fischer et al., 2016; Rieger & Massen, 2014; Wohlschläger, 2001), and has been hypothesized to play a role in both tool-object and body-tool interactions for a similar reason (Hegarty, 2004; Hegarty et al., 2005; Lesourd et al., 2025; Schwartz & Black, 1999; Schwartz & Holton, 2000; Wexler et al., 1998). It has been shown to be impaired in people with apraxia (Buxbaum et al., 2005; C. Evans et al., 2016; Ochipa et al., 1997; Roy et al., 1993; Rumiati et al., 2001; Schwoebel et al., 2004; Sirigu et al., 1996; Tomasino et al., 2003). We also tested the ability to mentally rotate a non-body object in space, which is often presumed to be required for performing both mechanical reasoning and tool selection tasks. In addition, we examined general reasoning and spatial working memory. Both of these processes have been implicated in mechanical reasoning (Cronbach, 1970; Hegarty, 2004; Hegarty et al., 1988; Mitko et al., 2024; Mitko & Fischer, 2020), although their relationships to tool use are less clear. Nevertheless, it is plausible that tool use could involve general reasoning to assess the compatibility of tool-object features or discern the underlying rules governing their use, or spatial working memory to track spatial relationships between tools, limbs, and recipient objects. By jointly examining whether these processes show similar patterns of association to mechanical reasoning, novel tool use, and limb apraxia, we aimed to provide further evidence of the extent to which mechanical reasoning is related to novel tool-use ability.

## Methods

### Participants

Thirty-one right-handed participants with chronic left hemisphere stroke (age = 63.03 ± 10.38; 12 Females; post-stroke months = 133.06 ± 70.76) were recruited. As part of a prior study, these participants completed the comprehension subtest of the Western Aphasia Battery (WAB) (Kertesz, 2007); all scored at least 4 out of 10 (moderate impairment). Fifteen neurotypical controls matched in age (age = 64 ± 10.14; 8 Females) and education (year = 14.77 ± 2.95 vs.15.47 ± 2.23) also participated in the experimental sessions. All participants gave written informed consent before participating, and the study was approved by the Institutional Review Board of Thomas Jefferson University. Participants were paid for their participation.

### Tasks and procedure

#### Stroke, Aphasia, and Apraxia testing

Participants completed a session of background testing to assess stroke and apraxia severity. The National Institutes of Health Stroke Scale was used to quantify general stroke severity in all individuals with stroke. Praxis ability was assessed using two tests, pantomime of tool use movements and imitation of meaningless movements, in all patients and 10 out of 15 healthy controls. Pantomimed tool use is a commonly used assessment (Goldenberg, 1996; Randerath et al., 2011) that is known to be highly correlated with actual tool use (Belanger et al., 1994; Clark et al., 1994; Hermsdörfer et al., 2013; Poizner et al., 1995) but is more sensitive to subtle tool-related impairments in people with apraxia as there are no cues or tactile feedback from the tool or recipient objects available to support grasp configuration and movement execution. Pantomimed tool use was assessed with a gesture-to-sight task, in which patients were shown a picture of a tool and asked to demonstrate how they would use that object (Watson & Buxbaum, 2015). Meaningless imitation was assessed by showing individuals a video of an actor performing an abstract movement and reproducing the same gesture in a mirror-like fashion (Buxbaum, Kyle, & Menon, 2005). For both tasks, movements were performed using the left (less affected) arm. Movements were video recorded and coded by two trained raters who achieved inter-rater reliability (Cohen’s Kappa >85%) on a subset of participants. Performance was scored based on three types of errors: incorrect hand posture, incorrect arm posture, and incorrect movement amplitude or timing, then summed to give a total score for each task.

As in our prior published research, we used the average score of the two tasks to classify all patients as apraxic or not (Buxbaum & Saffran, 2002): patients scoring 2 standard deviations below the mean neurotypical control score were classified as exhibiting apraxia. This yielded 15 and 16 participants in the non-apraxic and apraxic groups, respectively (Figure 1A). In recognition of the fact that individuals who were classed as apraxic or non-apraxic exhibited a range of scores on these two tests, we also conducted analyses that treated the scores as continuous measures of apraxia severity (see Data Analysis). Moreover, because the gesture-to-sight and meaningless imitation tasks, although often highly correlated (ρ = 0.65 in our sample, Figure 1A), probe different aspects of the apraxia syndrome (Buxbaum & Saffran, 2002), in these correlational analyses we treated the scores as two separate measures to see if any differences between them emerged.

**Figure 1:**
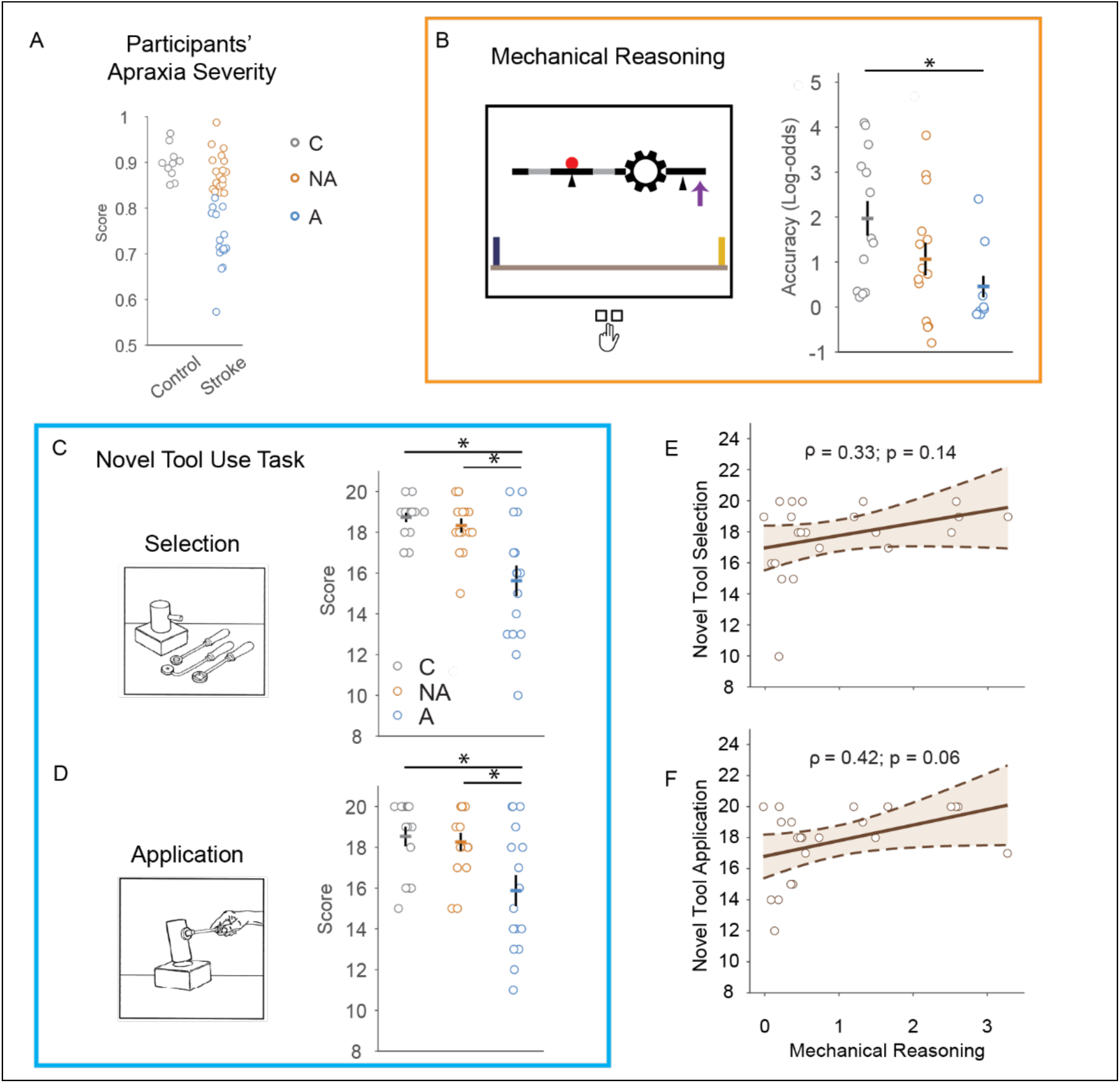
Performance in mechanical reasoning and novel tool use tasks. **A)** Individuals with left hemisphere stroke (n = 31) and matched neurotypical controls participated in our study (n = 15). Apraxia severity was assessed with pantomime and meaningless imitation. Using the average of these two scores, participants with left hemisphere stroke were divided into apraxic (A) and non-apraxic (NA) groups based on the performance of control (C) participants. **B)** We used a customized Rube Goldberg task to assess mechanical reasoning ability. Participants determined whether a mechanical system, based on the starting direction indicated by a purple arrow, would cause the red ball to roll toward the blue or yellow wall when it fell on the floor. Among participants who were able to complete the task (n = 35), response accuracy was different across groups. Specifically, the apraxic group had a lower accuracy than the control group, but not the non-apraxic group at step length of 3 (i.e., number of reasoning steps involved). There were no differences among groups at longer step lengths (i.e., 4–7; Supplemental Results) **C) & D)** Participants (n = 46) performed the novel tool use task created by Goldenberg & Hagmann (1998). The apraxic group had lower scores in both tool selection (C) and tool application (D) compared to the non-apraxic group and the control group, while no differences between the latter two groups were observed. **E) & F)** There were no significant correlations observed between mechanical reasoning and novel tool use, in particularly novel tool selection (E).

Participants completed the remaining experimental tasks in a second session, except the Corsi test (see below) that was completed with the praxis tasks. Each session lasted approximately 2 hours.

#### The Rube Goldberg task (Mechanical reasoning)

A Rube Goldberg machine, named after the famous cartoonist, consists of a chain of mechanical components that sequentially act on one another to achieve a task goal. We customized a Rube Goldberg task using four mechanical components: ramps, gears, levers, and pulleys. These common mechanical parts are frequently used to test mechanical aptitude in both industrial and scientific settings (Hegarty, 1992, 2004; Hegarty et al., 1988, 2005; Rozenblit et al., 2002; Schwartz & Black, 1996a, 1996b; Yoon & Narayanan, 2004). The task (as well as the mental rotation tasks described below) was coded using PsychoPy software (Peirce et al., 2019).

To perform the task, participants sat in front of a computer monitor. A static image of a Rube Goldberg machine was displayed at the center of the monitor (Figure 1B), with a thick gray bar appearing at the bottom of the screen (“the floor”). A short blue wall was presented at the left end of the floor and a yellow wall at the right end. A red ball was located somewhere within the machine. Participants needed to determine whether the mechanical system, based on the starting direction indicated by a purple arrow, would cause the red ball to roll toward the blue or yellow wall when it fell on the floor.

The task started with three screening/training blocks, each consisting of a demonstration and a criterion testing phase. The goal of these screening/training blocks was to ensure that people understood the basic physics of the individual components in isolation, before we tested their ability to reason about the components’ mechanical interactions. In the demonstration phase, participants were first given interactive instructions about how each mechanical component works using a question-and-answer format with the experimenter. For example, the experimenter presented an animation of the lever on the screen to participants and said “This is a lever. The lever can pivot like a seesaw: if one side goes down, the other side will go up. (As this is being said, the experimenter demonstrated the lever motion with their hands, pointing down and up respectively.) If a ball is placed at the center of the lever and this [left] side of the lever goes down, which way would the ball roll?” After responding, people were shown an animation of the lever moving and the resulting ball motion. Ramps and levers were introduced in training block 1, gears in training block 2, and pulleys in training block 3. There was no time limit for the demonstration, and explanations/animations could be repeated until the participant indicated that they understood how the mechanical component worked. Following the demonstration phase in each block, participants then completed a criterion test (5 to 7 trials) to determine whether they understood the functioning of each mechanical component. Participants were shown the component learned during the demonstration phase alone or paired with a previously learned component, and verbally or gesturally informed the experimenter which wall they thought the red ball would roll toward when it fell on the floor. Participants had to get 75% of criterion trials correct to advance to the next demonstration block. If they failed to reach 75% accuracy, they repeated the demonstration phase and criterion test. If they failed three rounds of demonstration and criterion testing for a given training block, they did not proceed to the main experimental test of the Rube Goldberg task. Two out of 15 participants in the neurotypical control group, one out of 15 participants in the non-apraxic group, and eight out of 16 participants in the apraxic group failed to pass the criterion trials. As these individuals struggled to understand the physics of individual components, we were unable to evaluate their ability to reason about mechanical interactions between these components. To check whether excluding these individuals might obscure a relationship between apraxia and mechanical reasoning, we simulated chance-level performance in the Rube Goldberg task for these individuals and noted that this did not qualitatively change our findings (Supplemental Figures S4 & S12).

Once participants passed all three training blocks, they moved on to a practice block and then two testing blocks. In these practice and test blocks, participants rested the index and middle fingers of their left hand on two keys (i.e., the z and x keys on a QWERTY keyboard). These two keys were marked by blue and yellow tape, respectively, that were spatially congruent with the blue and yellow walls on the screen. They were instructed to press the blue key if they thought the red ball would roll toward the blue wall and press the yellow key if the red ball would roll toward the yellow wall.

The practice block allowed individuals to practice the task and become familiar with using the response keys. There were 12 practice trials. Each trial started with a fixation cross presented at the center of the screen. After 1000 ms, the fixation cross was replaced by a Rube Goldberg machine, which remained on the screen for 60 s. If participants made a key press within 60 s, they received feedback about the accuracy of their response. A green check was displayed on the screen for 500 ms along with a pleasant auditory tone for a correct response, while a red cross and a buzzer tone occurred for an incorrect choice. People were given an error message if they took more than 60 s to respond. Following this feedback, the next trial then started after a 500 ms inter-trial interval. After the practice block, participants completed two testing blocks of 40 trials each. Testing blocks had the same setup as the practice block except that no feedback was given after each trial to avoid any learning during the test. Participants failed to respond on only 0.7 ± 0.2% of all trials (mean ± s.e.m.); thus, participants were typically able to make a response within 60 s. For each trial, a different Rube Goldberg machine with a varied number of mechanical components was displayed. The step length of each machine indicates how many mechanical components participants needed to consider (i.e., each component’s motion and interactions with other components) before determining the direction of the red ball when it hit the floor. For example, a lever acting on a gear would be a step length of 2: the first step determines how the ends of the lever move, and the second step determines the direction in which the gear rotates as a result of the lever’s action on it. Longer step lengths resulted in more difficult trials. Step lengths ranged from three to seven, with eight trials presented at each step length in each test block. Task performance was measured according to response accuracy and reaction time.

#### Novel Tool-Use Task

The novel tool use task, created by Goldenberg and Hagmann (Goldenberg & Hagmann, 1998), is frequently used to test participants’ ability to select and apply unfamiliar tools (Figure 2C). The test consisted of 10 trials. On each trial, participants were presented with 3 tools and 1 cylindrical object resting vertically on a base. In the selection phase, participants were asked to identify (without handling the tools) the tool that would best achieve the task of lifting the cylinder via a hole or “eye” in/on the cylinder. Participants received a score of 2 for selecting the correct tool, and lost 1 point for each wrong choice. Once the correct tool was identified, participants then performed the application phase in which they used the tool with their ipsilesional hand to lift the cylinder off the table, pause briefly (to demonstrate that the cylinder wouldn’t fall off the tool), then place the cylinder back on the table. Participants earned 2 points for successfully using the tool, but lost points for having difficulty applying the tool to the cylinder or dropping the cylinder while lifting it. The minimum score people could earn was 0 points in each phase of the task. Each participant was given a score, up to 20, for tool selection and tool application separately. All participants completed the novel tool use task.

**Figure 2:**
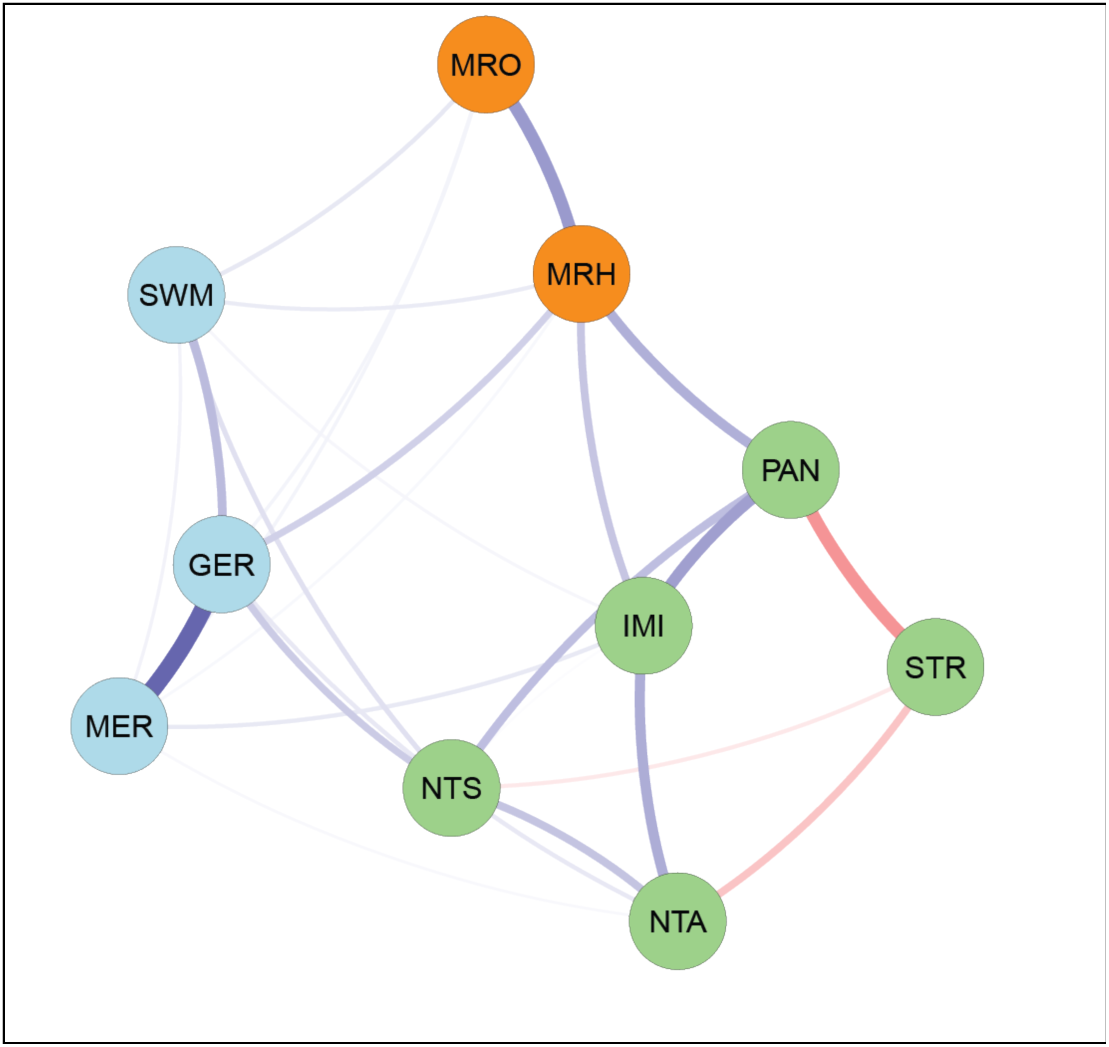
Latent structure among all task scores. A sparse network among variables established through the graphical lasso analysis. Using the Walktrap community detection algorithm applied to the network estimated via graphical lasso, we identified three distinct clusters of tasks (green, orange, and blue circles). Similar network structure was observed when pantomime tool use and meaningless imitation scores were combined as a single measure of apraxia (Figure S6). Thickness of lines: magnitude of partial correlation. Blue lines: positive partial correlation; Red lines: negative partial correlation. PAN: Pantomime tool-use; IMI: Meaningless Imitation; NTS: Novel tool selection; STR: NIH Stroke scale; NTA: Novel tool application; MRO: Mental rotation of object; MRH: Mental rotation of hand; SWM: Spatial working memory; GER: General reasoning; MER: Mechanical Reasoning.

#### Mental Rotation of Hand Task (Motor imagery)

Participants completed a task requiring them to mentally rotate images of hands which has often been proposed to reflect motor imagery (McAvinue & Robertson, 2008; Parsons, 1994; but see Vannuscorps et al., 2012). Participants sat in front a computer screen with their left index and middle fingers resting on the z and x keys. Their other hand rested on or below the desk (e.g., on their right leg) with the palm facing down toward the floor. In a familiarization block, participants were introduced to images representing either a right hand or a left hand. Each hand was presented from two perspectives, palm up or palm down. Participants then completed 8 trials in which they indicated with a key press whether the presented image was that of a right hand or a left hand. For this initial familiarization phase, each hand image was oriented with the fingertips pointing upward (i.e., not rotated). Each trial started with a fixation period for 1 s, followed by presentation of the hand stimulus. The trial ended upon a response or, if no response was made, 20 s after the image was presented. For this familiarization phase, participants received feedback indicating if their choice was correct. Participants proceeded to the next block if the response accuracy was not lower than 75% (6 out of 8 trials). In total, two out of 16 participants in the apraxic group and two out of 15 participants in the non-apraxic group failed to pass the criterion trials.

Participants next completed a practice block of eight trials. In the practice block, hand images were rotated at either 45 degrees or 225 degrees in the plane of the palm. This was followed by two testing blocks of 48 trials each, in which hand images were presented at rotation angles of 0, 60, 120, 180, 240, and 300 degrees (Figure S1A); each of the four hand images was presented twice at each rotation angle in randomized order. Participants were instructed to press a key with their left hand to indicate if the hand image was a right hand or a left hand as quickly and accurately as possible, and no feedback was provided during the testing blocks. Participants were required to respond within the 20 s limit (missed responses occurred on only 1.4 ± 0.5% trials). Reaction time and response accuracy were recorded.

#### Mental Rotation of Objects Task (Mental rotation)

This task was intended to assess participant’s ability to mentally manipulate non-body objects. In this standard 2D mental rotation task, participants were presented with pairs of arbitrary geometric shapes (Figure S1D) and were asked to determine whether the two shapes were the same or different. Here, “same” meant that the shapes were identical aside from a possible rotation in the plane of the screen, whereas “different” meant that the two shapes were mirror images of each other and thus could not be superimposed following a simple rotation in the plane of the screen (Cooper, 1975). In the familiarization block of eight trials, participants learned to indicate using key presses if the two presented shapes were the same or different. For this familiarization block, the two shapes were not rotated relative to each other.

If participants got at least 75% of the familiarization trials correct (four out of 16 participants in the apraxia group failed to pass this criterion), they next completed a practice block of eight trials. In each trial, the image on the right side of the screen was rotated by either 45 or 225 degrees relative to the image on the left. The shapes used in the practice block were different from those used in the familiarization and test blocks. In two subsequent testing blocks of 48 trials each, the relative rotation angle between the two images was either 0, 60, 120, 180, 240, 300 degrees, presented in randomized order. Participants had 20 s to press a key with their left hand (people failed to respond in only 2.2 ± 0.5% of all trials). Participants were instructed to respond as quickly and accurately as possible, and no accuracy feedback was provided during the testing blocks. Reaction time and response accuracy were recorded.

#### Raven’s Progressive Matrices, Short Form (General reasoning)

General reasoning ability was measured using a short version of the Raven’s Standard Progressive Matrices test (Figure S1E), which has previously been shown to correlate with the original 60-question test of reasoning ability (Bilker et al., 2012). People completed eighteen questions (i.e., both Form A and Form B, to increase reliability) using the same instructions and procedures as the standard version of the task. Briefly, on each trial participants were presented with an array of shapes, one of which was missing. People had to discern the underlying rule governing the visual relationship between the displayed shapes, then choose the correct option that would complete the array. The number of questions that participants correctly answered was used as the score. Only one participant from the apraxic group was unable to complete the task.

#### Corsi Test (Spatial working memory)

The Corsi test (Corsi & Michael, 1972) assessed visuo-spatial memory span. This task was administered in PEBL (Mueller & Piper, 2014). Individuals were presented with a scattered array of rectangles on a computer screen (Figure S1F). On each trial, rectangles would momentarily light up one by one in a particular sequence. After the sequence concluded, participants were required to use the computer mouse to click the rectangles in the same order that they were presented. The sequence length started at 2 rectangles and increased by 1 each time the sequence was correctly reproduced; participants had 2 attempts to reproduce a sequence of a given length. The task ended when the participant failed both attempts; the longest step length successfully completed was reported as the task score. Ten participants from the neurotypical group and 15 participants from each of the non-apraxic and apraxic groups completed this task.

### Data analysis

All statistical analyses were conducted in R (R Core Team, 2013). For the novel tool use task, the Corsi test, and the Raven’s matrix test, we analyzed the scores using one-way ANOVA (function: lm()) with participant group (neurotypical control, apraxia, non-apraxia) as the independent variable. For the Rube Goldberg (mechanical reasoning), mental rotation of hand (motor imagery), and mental rotation of objects tasks, we measured the response time and the binary response outcome (correct or incorrect) of each trial for each individual participant. We used a generalized linear mixed-effect model with a binomial link function to analyze binary response choices (function: glmer()). Fixed effects included the categorical variable group and continuous variable angle (in the mental rotation tasks) or step length (in the Rube Goldberg task); the models also included their interaction. In the mental rotation tasks, due to the rotational symmetry between 60 degree and 300 degrees, as well as between 120 and 240 degrees, the analysis was conducted using distinct angles 0, 60, 120, and 180 degrees. For the mental rotation of hand task, a fixed effect of hand posture of presented images (palm up or palm down) was also included. To account for within-subject variability, we included a random intercept and a random slope for angle/step length for each participant. Model comparisons based on the log-likelihood ratio test revealed that this model was favored over the model with only the random intercept or the random slope effect. For the response time data (correct trials only), we used a linear mixed-effect model that included the same fixed and random effects structure as the response choice model (function: lmer()). The random effect structure was also confirmed by model comparison. If necessary, post-hoc comparisons (function: emmeans()) were conducted, with correction for multiple comparisons using the Tukey method. The 95% confidence interval and effect size were also calculated (functions: confint() and eff_size()). When reporting results, we included main statistics including the 95% confidence interval and effect size. Note that effect size was not reported separately for the generalized linear mixed-effect models, as the fixed effect estimates themselves represent the effect size on the log-odds scale. In all analyses, the significance level was set at α = 0.05.

To address our second question of whether novel tool use and mechanical reasoning are associated with similar cognitive processes, and to complement analyses based on an arbitrary cutoff between participants with and without apraxia, we examined the correlation among task performance scores in all individuals with left hemisphere stroke by treating the severity of apraxia (measured by pantomime and meaningless imitation scores) as continuous measures. We also treated these scores as separate indices as they may reflect two distinct components of the apraxia syndrome (Buxbaum et al., 2014). Treating the average of two apraxia measures as a single apraxia severity score did not change the overall results (Supplemental Figures S5 & S6). The scores in the Rube Goldberg and two mental rotation tasks were calculated using the random coefficients from a generalized linear mixed-effect model on the binary response outcome data. Note the estimation of random coefficients in our original model introduced above was affected by the marginal mean (the mean among all three groups including the neurotypical control participants) since the random effect was assumed to be normally distributed around the marginal mean in mixed-effect models. Therefore, to more accurately estimate individuals’ scores for participants with left hemisphere stroke only, we re-ran the generalized linear mixed-effect model for these individuals without grouping (i.e., the only independent variable is angle/step length). Our results of the correlation analysis and the graphic analysis (described below) remained qualitatively unchanged by using the scores estimated from the original mixed-effect model that included all three participant groups.

Since the models can estimate the accuracy for all possible angles or step lengths, there are multiple scores for each individual in each task. We selected a single score for each task based on whether an interaction effect between group and rotation angle/step length was present in our original generalized linear mixed-effect models. For example, because of the significant interaction between step length and group in the Rube Goldberg task (see Results), accuracy was different at a step length of 3 among groups and this difference diminished at longer step lengths. Therefore, accuracy at longer step lengths may reflect an overall task difficulty effect that was not differentiable among groups, whereas accuracy at a step length of 3 better reflected the performance differences among the three groups. We thus used the accuracy estimated from the model for step length 3 in the correlation analysis. In contrast, for the mental rotation of hand and object tasks, no significant interaction between rotation angle and group was found (see Results), suggesting that performance difference among groups did not change with rotation angles. The score for each individual was, therefore, the estimated marginal response accuracy (i.e., mean across rotation angles). We confirmed that the results remained qualitatively unchanged if we used data at other rotation angles (e.g., 120 degrees).

To explore conditional dependencies and the latent structures among all tasks, we estimated a sparse partial correlation network on task performance using the Graphical LASSO algorithm (Friedman et al., 2008), which controls for spurious correlations by penalizing weak associations, yielding a more interpretable network for direct relationships. In particular, network estimation was performed using the EGA() function from the EGAnet package (Golino & Epskamp, 2017).

The Walktrap algorithm was used within the EGA() function to detect communities by identifying clusters of connected nodes that may reflect distinct latent processes.

Given the relatively modest sample size we had in this study, there is a concern that the resulting network might be unstable and that non-zero edges in the resulting network arise due to noise. We thus assessed the stability of the network structure through nonparametric bootstrapping (n = 1000) by resampling participants with replacement from the original dataset. Besides bootstrapping, because we used the estimated coefficients from regression models in this network analysis and these estimated coefficients had uncertainties (i.e., standard errors), we also assessed network stability by simulating synthetic data (n = 1000) using individual-level parameter estimates (i.e., mean and standard error of random intercept and random slope). We used both the bootstrapped samples and the simulated samples to assess network reliability. In particular, to assess the reliability of community detection, we evaluated the consistency with which each pair of tasks was assigned to the same community across 1000 bootstraps and across 1000 stimulated samples. To estimate edge-level stability, we estimated connection stability between each pair of tasks by calculating the proportion of bootstrapped and simulated networks in which each connection survived through the lasso regression.

### Imaging analysis

Structural T1 MRI scans were acquired from 27 of the 31 participants with left hemisphere stroke and used to derive lesion maps. Lesion maps were either hand-drawn or drawn using an automated algorithm (LINDA; Pustina et al., 2016), checked by an experienced neurologist (H. Branch Coslett) who was blinded to the goals of the study, then registered into the Montreal Neurological Institute (MNI) space (ch2bet template). Using these lesion maps, exploratory analyses aimed to identify lesions associated with impairments of mechanical reasoning, novel tool-use, and praxis (i.e., pantomime and meaningless imitation), as well as regions associated with the overlap of these impairments. To do this, we performed Support Vector Regression-Lesion Symptom Mapping (SVR-LSM) analyses using the SVR-LSM Matlab toolbox (DeMarco and Turkeltaub, 2018). SVR-LSM is a machine-learning based multivariate analysis technique that identifies relationships between lesioned voxels and behavior across all voxels and participants simultaneously. For mechanical reasoning, we conducted the analysis on the 19 participants with a lesion map available that passed the criterion in the Rube Goldberg task. The analyses for novel tool use and praxis were conducted with 27 participants. Only voxels lesioned in at least three participants were included in the respective analysis (see Figures 5A & B). Total lesion volume was regressed from both behavioral scores and the lesion maps. Voxelwise statistical significance was determined using a Monte Carlo style permutation analysis (10,000 iterations) in which the association between behavioral data and lesion map was randomized. A voxelwise threshold of p < 0.01 and a minimum cluster size of 100 voxels was applied to identify significant lesion-symptom relationships. SVR-LSM analyses were run on scores from the Rube Goldberg task, the selection score from the Novel Tool Use Task, and the overall Apraxia score (average score from the pantomimed tool use and meaningless imitation tasks). We then overlaid the results of these three analyses and looked for regions of overlap. Localization of significant grey and white matter clusters were identified with the Eve Atlas (John Hopkins University; Oishi et al., 2009); https://github.com/muschellij2/Eve_Atlas).

## Results

### Question 1: Is mechanical reasoning impaired in apraxia and is it related to novel tool use?

To examine the relationship between mechanical reasoning and novel tool use, in particular tool selection, we independently evaluated mechanical reasoning with the Rube Goldberg task and tool-use ability with the classic novel tool use task. Individuals with left hemisphere stroke were divided into two groups based on their overall apraxia severity (average of performance on the gesture-to-sight and meaningless imitation tasks, relative to neurotypical controls; see Methods). This resulted in 15 participants in the non-apraxic group and 16 participants in the apraxic group, along with a group of 15 neurotypical controls (Figure 1A).

#### Mechanical reasoning ability was similar for individuals with stroke with and without apraxia

Mechanical reasoning was tested using our Rube Goldberg task (Figure 1B). For participants who passed the criterion blocks and proceeded to the experimental assessment (n = 35; see Methods), we primarily focused on their response accuracy, as this reflects the overall capacity to reason about physical interactions between objects. A generalized mixed effect model revealed a significant effect of group on response accuracy, as well as an interaction between group and task difficulty (i.e., step length; See Supplemental Results & Supplemental Figure S2). In particular, when the task required three steps of reasoning (e.g., three mechanical components), the apraxic group was significantly less accurate than the control group (mean difference = −1.55 ± 0.62, p = 0.03, 95% CI = [−3.01, −0.08]; Figure 2B). However, there was no significant difference between either the apraxic and non-apraxic groups (mean difference = −0.87 ± 0.60, p = 0.32, 95% CI = [-2.28, 0.55]) or between the non-apraxic and control groups (mean difference = −0.68 ± 0.54, p = 0.42, 95% CI = [-1.94, 0.59]). At larger step lengths (i.e., step lengths 4-7), none of the three groups were significantly different from each other (all p > 0.08). These results did not change when we included simulated data from the remaining participants who did not pass the criterion tests (see Supplemental Figure S4).

In terms of response times, we did not observe an effect of group (F_2, 32.52_ = 2.46, p = 0.10) nor an interaction between group and step length (F_2, 32.22_ = 2.01, p = 0.15; Supplemental Figure S3A). This suggests that observed differences in response accuracy were unlikely to arise from a speed-accuracy trade-off.

#### People with apraxia were particularly impaired at selecting and applying novel tools

All participants (n = 46) completed the novel tool use task. Participants first selected the tool that would allow them to complete the mechanical problem (i.e., lift the cylinder from its base; Figure 1E). We observed a significant effect of group on tool selection performance (F_2,43_ = 11.31, p = 0.0001). The apraxic group had a lower tool-selection score than both the control (mean difference = −3.11 ± 0.72, p = 0.0002, 95% CI = [−4.84, −1.37], effect size= −1.56) and the non-apraxic groups (mean difference = −2.71 ± 0.72, p = 0.001, 95% CI = [−4.44, −0.97], effect size = - 1.36). In contrast, no significant difference was observed between the non-apraxic and control groups (mean difference = −0.40 ± 0.73, p = 0.85, 95% CI = [-2.41, 1.36], effect size = −0.20).

Participants were then asked to physically apply the correct tool to solve the mechanical problem (e.g., lift the cylinder from its base using the ipsilesional hand). Beyond the cognitive processes required in tool selection, physically applying a tool also involves motor planning and execution. There was a significant effect of group on the ability to successfully use the novel tools (F_2,43_ = 6.37, p = 0.0037; Figure 1F). In particular, performance was worse in the apraxic group compared to both the control group (mean difference = −2.66 ± 0.83, p = 0.0067, 95% CI = [−4.66, −0.66], effect size = −1.16) and the non-apraxic group (mean difference = −2.39 ± 0.83, p = 0.016, 95% CI = [−4.39, −0.39], effect size = −1.04). Again, we observed no difference between the control and non-apraxic groups (mean difference = −0.27 ± 0.84, p = 0.95, 95% CI = [-2.32, 1.77], effect size = −0.12). These results echo prior reports (Goldenberg & Hagmann, 1998; Jarry et al., 2013; Osiurak et al., 2013) in highlighting the impact of limb apraxia on the ability to properly select and apply novel tools.

#### Mechanical reasoning ability was not correlated with novel tool use ability

Since mechanical reasoning and performance on the novel tool-use task were both impaired in individuals with limb apraxia relative to controls, we tested whether the same individuals were likely to exhibit impairments on these two tasks. That is, we examined the correlation between mechanical reasoning (as measured at a step length of 3) and the accuracy of either selecting or applying tools in the novel tool-use task. For this analysis, we included individuals with left hemisphere stroke regardless of apraxia severity. We observed no significant correlations between either tool selection or tool application and mechanical reasoning (selection: ρ = 0.33, p = 0.14; application: ρ = 0.42, p = 0.06; Figure 1E-F; Supplemental Figures S5, S6). Importantly, a number of participants demonstrated marked impairments in mechanical reasoning but achieved ceiling or near-ceiling performance in the novel tool use task. These results thus suggest that even though people with apraxia tend to be impaired relative to the neurotypical control group in each of these two tasks, the ability to perform one task is not strongly indicative of the ability to perform the other.

### Question 2: Are the same cognitive processes associated with mechanical reasoning and tool use?

A second way to examine the hypothesis that mechanical reasoning is not strongly related to novel tool selection or application is to assess whether mechanical reasoning and novel tool use abilities rely on (or at least bear a similar relationship to) the same cognitive processes. To determine this, participants completed four additional tasks that have previously been proposed to be related to mechanical reasoning and/or tool-use abilities in the literature: motor imagery (mental rotation of hand task), mental rotation (mental rotation of object task), general reasoning (Raven’s matrices task), and spatial working memory (Corsi test). Performance in these tasks exhibited different patterns among participant groups (See Supplementary Results; Supplemental Figure S1).

#### Mechanical reasoning is associated with other reasoning abilities but not with novel tool use

To examine how these different cognitive capacities relate to each other and to mechanical reasoning and novel tool use, we examined the correlational relationships across the performance of all tasks. Since performance on a given task may not solely depend on unique cognitive capacities, we focused on the conditional dependencies and latent clusters among tasks examined by graphic lasso regression that estimates partial correlations between each pair of tasks while controlling for all the others (Friedman et al., 2008). To account for the effect of overall stroke severity in these partial correlation estimates, we also included the NIH Stroke Scale score in these analyses.

The global organization of all tasks revealed that mechanical reasoning and novel tool selection/application belonged to two distinct latent clusters (Figure 2). Specifically, mechanical reasoning clustered with general reasoning and spatial working memory, reflecting a latent construct centered around high-level reasoning abilities. This cluster did not include novel tool selection or application. Indeed, no direct link was identified between mechanical reasoning and either novel tool selection or application in the sparse network, consistent with the absence of a zero-order correlation between them (Supplemental Figures S5-S6).

The separation between the clusters involving novel tool use and mechanical reasoning was found to be robust based on our bootstrap and simulation analyses (Supplemental Figures S8 & S10). In particular, 86.6% of the bootstrapped networks (n = 1000) and 95.7% of the simulated networks (n = 1000) exhibited at least two clusters, which in all cases separated novel tool selection and mechanical reasoning into distinct clusters (Supplemental Figures S8A & S10A). In many cases a third cluster was also identified, which included the two mental rotation tasks (see Supplementary Results of Sparse Network). Thus, mechanical reasoning was associated with a distinct latent factor and was reliably disconnected from novel tool selection.

#### Novel tool use associates with other praxis abilities

In contrast to mechanical reasoning, novel tool selection formed a separate latent cluster with other praxis-related abilities. For these analyses, performance on the pantomimed tool-use and meaningless imitation tasks were treated separately as they may reflect distinct aspects of the apraxia syndrome and have been shown to partially dissociate behaviorally and neurally in prior studies (Buxbaum, 2017; Rothi & Heilman, 2014). Using a single apraxia severity score as above did not change the overall results of these analyses (Supplemental Figure S6).

We noted that novel tool selection grouped with novel tool application, pantomime, and meaningless gesture imitation, which together represent the well-established trio of abilities (i.e., tool use, pantomime, and imitation) that are traditionally associated with praxis. Novel tool selection was consistently connected with pantomimed tool use but not with imitation across our original data, bootstrapped, and stimulated samples. Interestingly, while there was no connection between mechanical reasoning and pantomimed tool use, we noted a subtle but reliable connection between mechanical reasoning and meaningless gesture imitation (Supplemental Figures S7-S11).

We also did not observe evidence of a direct connection between novel tool selection or application and the mental rotation of either hands (i.e., motor imagery) or objects. Taken together, these results suggest that mechanical reasoning is not a primary, independent predictor of tool selection ability or overall praxis, and that novel tool selection is instead associated with a distinct latent praxis construct that is separable from other examined cognitive abilities.

#### Mechanical reasoning and tool use rely on distinct brain regions

Exploratory lesion-symptom mapping (SVR-LSM) analyses were conducted to assess the neural underpinnings associated with impairments to mechanical reasoning, novel tool selection, and praxis abilities (i.e., the average score of pantomime tool-use and meaningless imitation). Strikingly, we found that impairments in each of these three tasks are associated with lesions to distinct brain areas. We found that praxis impairments were primarily associated with lesions in the posterior middle temporal gyrus (pMTG) and inferior parietal lobule (IPL), including the posterior supramarginal gyrus (pSMG; p < 0.01; Figure 3C and D in red; Table 1A). Impairments in novel tool selection were mostly related to lesions impacting the postcentral gyrus and the IPL, including the anterior supramarginal gyrus (aSMG; p < 0.01; Figure 3C and D in yellow; Table 1C), consistent with previous findings (Goldenberg & Spatt, 2009; Orban & Caruana, 2014; Reynaud et al., 2016). The two maps, when superimposed, revealed a small 151-voxel area of overlap within the supramarginal gyrus (depicted in orange Figure 3C & D). Finally, mechanical reasoning impairments were linked to lesions in the precentral gyrus, inferior and middle frontal gyri (IFG and MFG), as well as the surrounding white matter including the corona radiata and external capsule (p < 0.01; Figure 3C and D in blue; Table 1B); no clusters overlapped the brain regions associated with either apraxia or novel tool selection.

**Figure 3:**
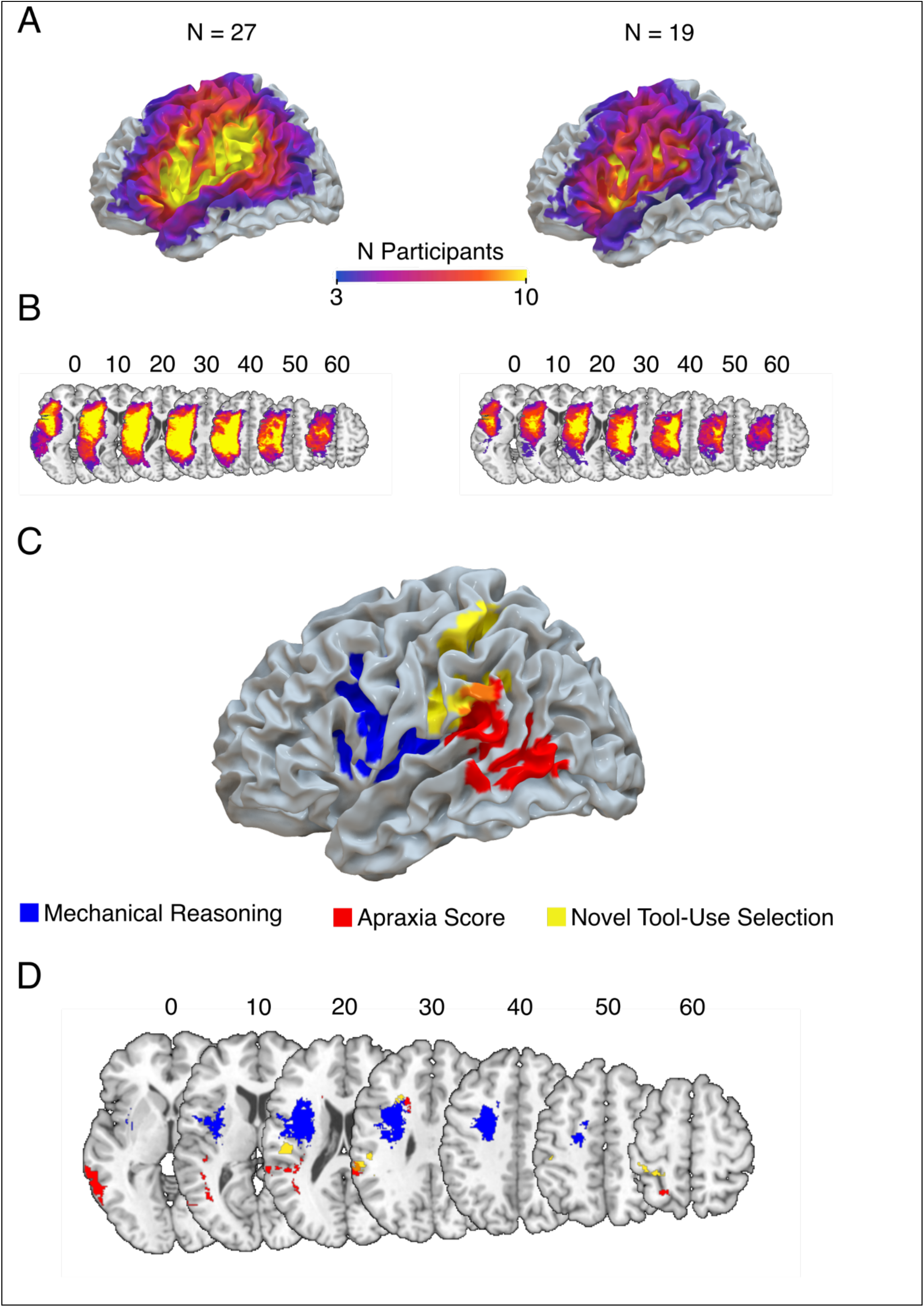
Lesion-symptom mapping. **A)** Lesion overlap maps overlaid on brain render. On left is the map for the 27 participants included in the analysis of praxis and novel tool-use abilities. On the right is the map for the 19 participants included in the analysis of the mechanical reasoning data. **B)** Lesion overlap maps represented on axial brain slices. **C)** Brain render with the lesion correlates associated with praxis abilities (average of pantomime and meaningless imitation) in red, novel tool-use selection in yellow and mechanical reasoning in blue. **D)** The lesion correlates for each of the three tasks are represented on axial brain slices. The overlap between praxis abilities and novel tool-use is represented in orange (slice z = 30, also visible on C).

**Table 1:** Neural correlates of impaired performance. A) Praxis. B) Mechanical reasoning. C) Novel tool-use selection. Coordinates of the cluster centroid are in MNI space.

| *Brain area(s)* | *Cluster extent*<br>*(mm<sup>3</sup>)* | *Peak Z-*<br>*value* | *Cluster centroid*<br>*coordinates* |
| --- | --- | --- | --- |
| **(A) Apraxia Score (Pantomime Tool-Use and Meaningless Imitation)** |  |  |  |
| Occipito-Temporal<br>*Middle Temporal Gyrus*<br>*Inferior Occipital Gyrus* | 2396 | -3.72 | -60; -54; 2 |
| Parieto-Temporal<br>*Supramarginal Gyrus*<br>*Superior Temporal Gyrus* | 1541 | -3.72 | -51; -38; 23 |
| Middle Temporal Gyrus | 708 | -3.54 | -41; -51; 14 |
| Corona Radiata<br>Corpus Callosum | 344 | -3.72 | -18; 17; 29 |
| Superior Parietal Lobule | 182 | -2.47 | -22; -57; 58 |
| **(B) Mechanical Reasoning** |  |  |  |
| Frontal (cortical and subcortical)<br>*Superior Corona Radiata*<br>*Precentral Gyrus*<br>*Inferior Frontal Gyrus*<br>*Middle Frontal Gyrus*<br>*External Capsule*<br>*Superior Frontal Gyrus*<br>*Superior Longitudinal Fasciculus*<br>*Insula*<br>*Postcentral Gyrus*<br>*Putamen* | 17,794 | -3.72 | -33; -1; 27 |
| **(C) Novel Tool Selection** |  |  |  |
| Inferior Parietal<br>*Supramarginal Gyrus*<br>*Postcentral Gyrus* | 1,300 | -3.54 | -53; -27; 26 |
| Superior Parietal<br>*Postcentral Gyrus*<br>*Superior Parietal Lobule* | 582 | -3.16 | -32; -40; 58 |
| Middle Frontal Gyrus | 121 | -2.85 | -26; 20; 31 |
| Postcentral Gyrus | 114 | -3.16 | -44; -28; 50 |
| Inferior Parietal<br>*Angular Gyrus*<br>*Supramarginal Gyrus* | 101 | -3.09 | -50; -44; 34 |

## Discussion

The ability to properly select and apply a novel tool to complete a desired task has long been found to be impaired in limb apraxia. However, the underlying factors contributing to this deficiency remain a matter of debate. In this study, we tested a contemporary hypothesis that novel tool use deficits (and tool-use deficits more broadly) in apraxia are caused by impaired mechanical reasoning – step-by-step, deliberative mental simulation of how one object causally affects another (Allen et al., 2020; Hegarty, 2004), as required in our Rube Goldberg task. Consistent with previous findings, we demonstrated that individuals with limb apraxia performed worse on a classic novel tool-use task than individuals without apraxia. However, when mechanical reasoning was measured independently from novel tool use ability, we found no evidence that it was specifically associated with apraxia or correlated with novel tool selection/application. Moreover, mechanical reasoning and novel tool use ability were found to be associated with distinct cognitive abilities and neural substrates. These findings suggest that the ability to select and apply the correct tool to perform a task is unlikely to rely on the ability to reason about the mechanical interactions between tools and objects.

Prior work has often used the novel tool use task, or closely-related variations, to measure mechanical reasoning ability (Baumard et al., 2014, 2016, 2021; Faye et al., 2018, 2018; Goldenberg & Hagmann, 1998; Hodges et al., 1999; Jarry et al., 2013; Osiurak et al., 2009; Seifert et al., 2025; Spatt et al., 2002). These studies have reported correlations between performance on novel tool use (purported to be a measure of mechanical reasoning) and measures of pantomimed tool use, meaningless imitation, and familiar tool use (Bartolo et al., 2007; Goldenberg, 2013a; Hartmann et al., 2005; Jarry et al., 2015; Osiurak et al., 2009). Based on those associations, it has been argued that mechanical reasoning is critical to tool use and that impairment of this ability underlies various manifestations of apraxia (Baumard et al., 2014, 2021; Goldenberg & Hagmann, 1998; Jarry et al., 2013, 2015; Osiurak et al., 2009; Osiurak, Reynaud, et al., 2021). However, as noted earlier, assessing mechanical reasoning through novel tool use tasks introduces a circular logic: impaired tool use is assumed to indicate impaired mechanical reasoning, which is then taken as a causal factor for any observed tool-use deficits. In contrast, a key strength of our study is the independent assessment of mechanical reasoning and novel tool use abilities. By disentangling these measures, we revealed a behavioral dissociation whereby individuals with apraxia exhibited greater deficits in novel tool selection and application but comparable mechanical reasoning performance to individuals with left hemisphere strokes without apraxia. Furthermore, our data did not support a direct correlation between mechanical reasoning and novel tool-use abilities. Instead, in line with prior hypotheses in the literature (Buxbaum 2017), these abilities were primarily associated with different aspects of praxis: novel tool selection was strongly correlated to pantomimed tool use, while there was a weak but reliable correlation between mechanical reasoning and the imitation of meaningless gestures – a task that does not involve any tool use. This dissociation was further reiterated by our exploratory neuroimaging data, in which we observed that more anterior brain regions were associated with mechanical reasoning whereas more posterior brain areas (including inferior parietal regions classically linked to apraxia) were associated with novel tool selection (Buxbaum et al., 2014; Buxbaum & Randerath, 2018; Goldenberg & Randerath, 2015; Goldenberg & Spatt, 2009; Orban & Caruana, 2014; Reynaud et al., 2016). Together, these results provide compelling evidence that mechanical reasoning is not essential for using novel tools.

One important concern is whether the dissociation observed between mechanical reasoning and novel tool selection or application simply reflects a difference in the specific kind of mechanical components involved in each task. If so, it could be argued that individuals with apraxia struggle specifically with tools but not with other kinds of mechanical or physical principles such as those featured in the Rube Goldberg task. However, many everyday tools and machines, such as can openers, pliers, crowbars, screws, and blinds and curtains, are fundamentally built on the same mechanical components used in the Rube Goldberg task (Hegarty, 1992, 2004; Hegarty et al., 1988, 2005; Mitko & Fischer, 2020; Rozenblit et al., 2002; Schwartz & Black, 1996a, 1996b; Yoon & Narayanan, 2004). These components also form the basis of standardized assessments of mechanical aptitude that are widely used in industrial settings (Corporation, 1990). Therefore, we argue that the Rube Goldberg task serves as a representative measure of the same kind of mechanical reasoning ability that would be most likely to be required by conventional tools and devices. Moreover, novel tools are simply arbitrary objects whose functions are yet to be defined or discovered by the user; people are more likely to attribute a basic set of simple mechanical principles to such objects including those we present in our Rube Goldberg task. The observed dissociation between this general form of mechanical reasoning and the ability to select and use novel tools (e.g., in both zero-order correlations and the graphical LASSO analyses) therefore suggests that novel tool use requires distinct cognitive processes from those engaged by mechanical reasoning.

Although our data suggest that mechanical reasoning is largely dissociable from apraxia, it is noteworthy that eight participants in the apraxic group did not pass the criterion test in the Rube Goldberg task. On one hand, this might suggest that mechanical reasoning may be impaired in highly apraxic individuals. However, our simulation analysis sought to address this concern by assuming that these participants would perform at chance if they had been tested on the Rube Goldberg task, since they were unable to demonstrate a basic understanding of the individual mechanical components in the criterion tests. These simulations did not reveal any evidence of a potential relationship between mechanical reasoning and novel tool selection despite the assumption that these individuals would perform more poorly than the other participants on the Rube Goldberg task. On the other hand, the fact that these participants failed to understand the individual mechanical components in the criterion tasks raises the possibility that some individuals with apraxia may struggle with an intuitive understanding of basic physical properties such as gravity, rotation, collisions, balance, and so on – hallmarks of so-called intuitive physics (Gilden & Proffitt, 1989; Krist et al., 1993; Kubricht et al., 2017; Mitko & Fischer, 2020). Under this hypothesis, impairments in the ability to draw inferences about basic physical properties, rather than in higher-level mechanical reasoning about interactions between objects, could contribute to the tool-use difficulties observed in apraxia. However, a closer inspection of our data does not support this possibility. A few neurotypical participants and participants with strokes without apraxia also failed the criterion test in the Rube Goldberg task, suggesting that the failure is not specific to apraxia. Furthermore, examination of the performance of the individuals who failed the Rube Goldberg criterion test indicated that their performance also tended to be lower in all other tasks (Supplemental Figure S12). This suggests that there is nothing particularly special about the poor performance of these individuals in the Rube Goldberg task: the ability to understand the physical properties of mechanical components is unlikely to be specifically indicative of novel tool use ability. Indeed, a recent case series in our lab revealed a dissociation between praxis and the ability to intuitively infer the basic physical properties of objects (Liu et al., 2025).

If mechanical reasoning is not the key factor, what else might explain impaired novel tool selection or application in apraxia? A popular hypothesis proposes that tool-use deficits arise from impaired simulation of how the body interacts with tools and objects (i.e., motor imagery), rather than between tools and objects absent the body (Buxbaum, 2017). This hypothesis proposes a kind of embodied simulation that is distinct from the mental simulation of body-independent interactions between objects (Amorim et al., 2006; Kosslyn et al., 1998; Reed et al., 2004). Regardless of whether such a form of embodied simulation exists (e.g., Vannuscorps et al., 2012), our results highlight a lack of a direct link between the motor imagery and novel tool use tasks, and sequester mental rotation of bodies and objects into a distinct latent cluster separate from praxis abilities. On the other hand, while there is no link to a task requiring the use of actual tools, we do see an association between this cluster and more abstracted forms of praxis (i.e., imitation and pantomimed tool use) in line with the prior literature (Buxbaum et al., 2005; C. Evans et al., 2016; Ochipa et al., 1997; Roy et al., 1993; Rumiati et al., 2001; Schwoebel et al., 2004; Sirigu et al., 1996; Tomasino et al., 2003). Overall, this suggests that the relationship between mental simulation writ large and praxis might be largely mediated by additional task demands, namely having to imagine how to interact with an object that is not present or when planning the use of tools that have fairly uninformative affordances (e.g., Thibault et al., 2025).

The absence of associations between novel tool selection and mechanical reasoning or motor imagery implies that in many cases, successfully selecting a tool is unlikely to depend on effortful deliberation or any form of mental simulation. Instead, we propose that novel tool selection relies more heavily on intuitive processes (i.e., System 1, (e.g., Evans & Over, 2013; Kahneman, 2011)), whereby a tool is chosen automatically according to existing action-object associations rather than through effortful, time-consuming reasoning. It is thought that we store abstract semantic representations of tools and their associated actions (e.g., action semantics; (van Elk et al., 2014)), which can be rapidly and automatically evoked to influence actions (Bub et al., 2008; Myung et al., 2010; Tucker & Ellis, 1998). In everyday novel tool contexts, we may simply extrapolate from these existing action-semantic representations to enable fast, intuitive tool selection without the need for explicit mechanical reasoning. For example, a representation of the swinging action associated with a tennis racket can be used to facilitate initial game play of other racket sports like squash or ping pong. This is strikingly different from deliberating how a squash racket works before its initial use – something that can be done, but may not be necessary and might even become counterproductive (Beilock et al., 2004; Fitts & Posner, 1967).

Novel tool use may also rely on other automatic processes such as the ability to rapidly perceive visual relations between objects (Hafri & Firestone, 2021). Our results revealed a correlation between novel tool selection and general reasoning, as measured by the Raven’s Matrices test. Although the Raven’s Matrices test is designed to primarily assess general reasoning ability (i.e., to identify the rule governing the relationship between a set of stimuli), it is plausible that this task is supported by a more automatic recognition of visual patterns (e.g., when scanning the set of available response options to identify which one matches an inferred solution). The classic novel tool use task (Goldenberg & Hagmann, 1998) appears to require a similar ability to match complementary shapes of tools and recipient objects, and this automatic relation perception is likely a key factor underlying success in selecting the correct tool in real life activities such as choosing a screwdriver tip to match a specific screw head – a chore that likely does not require mental simulation. Thus, novel tool selection and general reasoning ability may indeed be linked by shared underlying cognitive processes, but those processes need not necessarily require deliberative reasoning. Critically, while both mechanical reasoning and novel tool use are linked to general reasoning ability, they are not associated with each other, suggesting that the link to general reasoning arises from different underlying cognitive components of the Raven’s Matrices task. In other words, effective tool selection can occur without invoking resource-demanding processes such as mechanical reasoning – processes that may be unnecessarily invoked by asking people to explicitly think about tool use in laboratory settings.

In summary, our findings challenge the account that mechanical reasoning is a necessary and causal basis for the successful selection and use of novel tools. By independently measuring mechanical reasoning and novel tool use, we found no behavioral or neural evidence supporting a direct relationship between these two abilities. More broadly, our data suggest that tool use and mechanical reasoning are related to different clusters of cognitive abilities. Critically, at present it appears that the ability to select and apply novel tools is more closely aligned with other praxis abilities than with mechanical reasoning or other forms of high-level reasoning processes. Based on these findings, we propose a shift away from deliberation-centered accounts of tool use and toward models emphasizing intuitive and automatic tool selection.

## Funding

This work was supported by NIH Grant #R01NS115862 to Aaron L. Wong.

## Supplementary Results

### Motor imagery was impaired in individuals with apraxia

Data were analyzed from participants who passed the motor imagery criterion tests (n = 42). A main effect of rotation angle was observed (χ^2^_1_ = 59.72, p < 0.0001; Figure S1B), which did not significantly interact with group (χ^2^_2_ = 1.46, p = 0.48) or hand-image orientation (χ^2^_1_ = 1.04, p = 0.31). Specifically, response accuracy decreased with rotation angle (slope = −0.88 ± 0.11, 95% CI = [−1.11, −0.66]). We also observed effects of group (χ^2^_2_ = 19.04, p < 0.0001), hand-image orientation (χ^2^_1_ = 4.42, p = 0.036) and their interactions (χ^2^_2_ = 14.45, p = 0.0007). No significant three-way interaction was found (χ^2^_2_ = 0.53, p = 0.77). Post hoc comparisons revealed that response accuracy was lower for the palm-up images than for the palm-down images in the apraxic group (mean difference = −0.45 ± 0.13, p = 0.0005, 95% CI = [−0.70, −0.20]), but not in the non-apraxic group (mean difference = −0.33 ± 0.20, p = 0.079, 95% CI = [-0.71, 0.04]) or the control group (mean difference = 0.29 ± 0.18, p = 0.1012, 95% CI = [-0.06, 0.65]). We also compared the response choice performance between groups for the palm-up and palm-down orientation conditions, respectively (Figure S1C). When a palm-up image was shown with no rotation, response accuracy was lower in the apraxic group compared to the control group (mean difference = −1.79 ± 0.41, p < 0.0001, 95% CI = [−2.43, −0.46]) and the non-apraxic group (mean difference = −1.45 ± 0.42, p = 0.002, 95% CI = [−2.43, −0.46]). No difference was found between the non-apraxic and control groups (mean difference = −0.34 ± 0.43, p = 0.71, 95% CI = [-1.34, 0.66]). Similar results were observed for the palm-down images. The apraxic group had a lower accuracy than the control group (mean difference = −1.05 ± 0.41, p = 0.028, 95% CI = [−2.00, −0.09]) and the non-apraxic group (mean difference = −1.33 ± 0.43, p = 0.005, 95% CI = [−2.34, −0.33]). No significant difference was found between the non-apraxic group and the control group (mean difference = 0.29 ± 0.43, p = 0.78, 95% CI = [-0.72, 1.29]). Note that the lower accuracy in the apraxic group was not a result of a speed‒accuracy trade-off, as their response times were slower than those of the other two groups (Supplementary Figure S3C). Taken together, these results are consistent with the well-established finding that motor imagery is impaired in people with apraxia.

**Figure S1:**
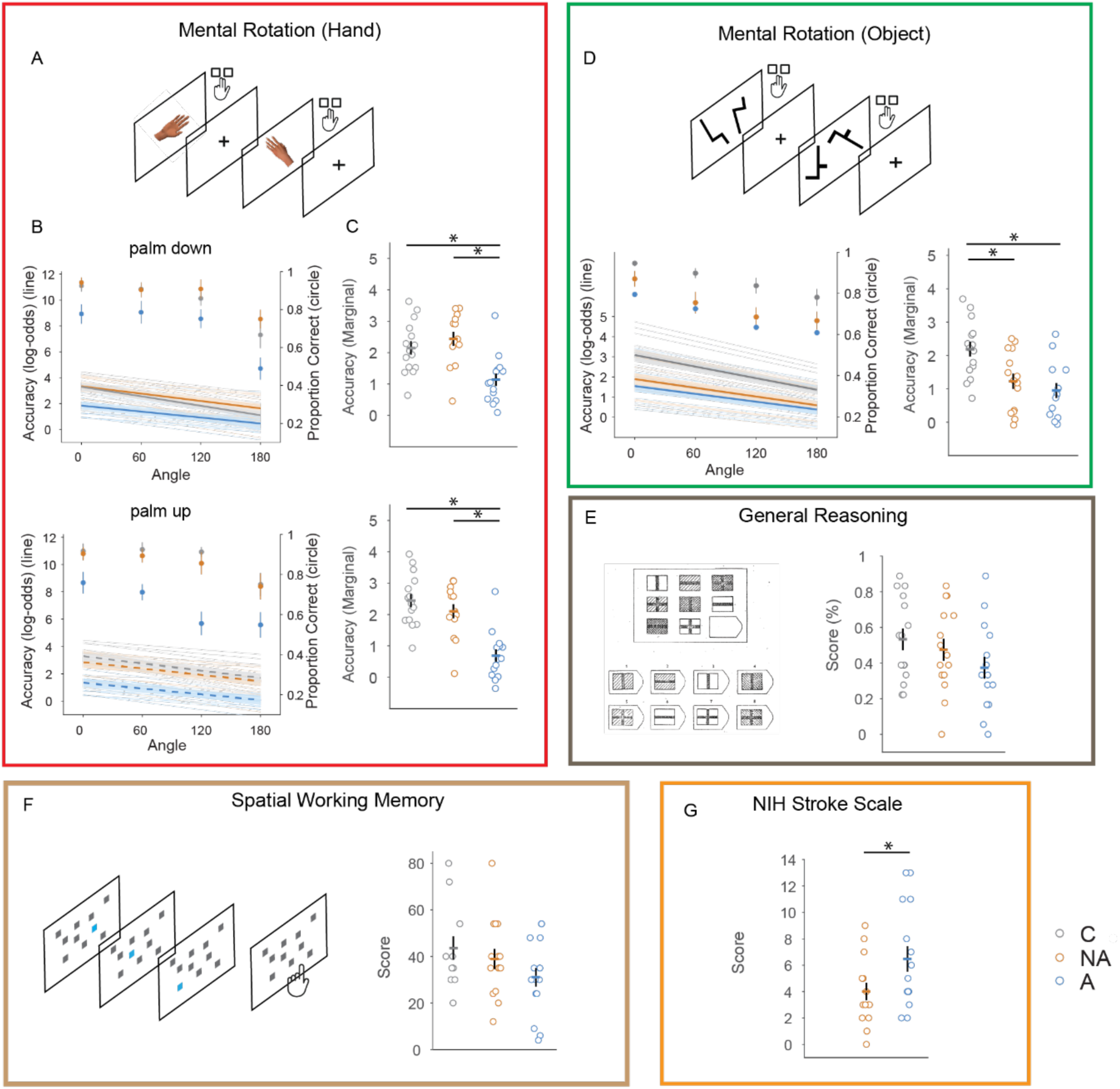
Performance of motor imagery, mental rotation (object), general reasoning, spatial working memory, and NIH stroke scale tests. **A)** Motor imagery was assessed by asking participants to determine the laterality of a rotated hand image. **B)** Response accuracy was affected by group, rotation angle, and hand image orientation (palm-down versus palm-up). **C)** Group comparisons revealed that the apraxic group had significantly lower accuracy than both the non-apraxic and control groups, regardless of hand image orientation. **D)** Top panel: In the mental rotation task, participants judged whether two shapes were identical or mirrored. Left panel: Accuracy varied across groups and declined with increasing rotation angles. Right panel: Post hoc analysis showed that both the apraxic and non-apraxic groups performed worse than the control group. **E)** General reasoning ability was measured using a short version of the Raven’s Progressive Matrices test. Although no statistical differences were observed, scores exhibited a decreasing trend from the control group to the non-apraxic group and then to the apraxic group. **F)** Spatial working memory was assessed by the Corsi task. Similar to general reasoning, no group differences were observed, yet a decreasing trend was visualized from the control group to the non-apraxic group and then to the apraxic group. **G)** Stroke severity was quantified by the NIH Stroke scale, administered only to patient groups. Scores were higher (more severe stroke) in the apraxic group compared to the non-apraxic group.

### Mental rotation of non-body objects was impaired in individuals with stroke

Response choice data from participants who passed the criterion test (n = 42) showed significant effects of group (χ^2^_2_ = 10.69, p = 0.0047) and rotation angle (χ^2^_1_ = 77.79, p < 0.0001), but no significant interaction between group and rotation angle (χ^2^_2_ = 2.48, p = 0.29) (Figure S1D, left panel). On average, response accuracy decreased with increasing rotation angles (slope = −0.78 ± 0.09, 95% CI = [−0.95, −0.61]). Post hoc group comparisons revealed that both the apraxic group (mean difference = −1.26 ± 0.38, p = 0.002, 95% CI = [−2.14, −0.39]) and the non-apraxic group (mean difference = −0.98 ± 0.36, p = 0.016, 95% CI = [−1.81, −0.15]) had worse performance than the control group. However, no difference was found between the apraxic and non-apraxic groups (mean difference = −0.28 ± 0.37, p = 0.72, 95% CI = [-1.14, 0.58]). No group difference (F_2, 39.11_ = 2.51, p = 0.094; Supplementary Figure S3C) or interaction between group and rotation angle (F_2, 39.06_ = 0.69, p = 0.51) were observed for response times. These results demonstrate that unlike motor imagery, mental rotation of objects is not specifically impaired in people with apraxia.

### General reasoning was not impaired in individuals with apraxia

Participants (n = 45) completed a short form of the Raven’s Progressive Matrices task. No significant group differences were observed (F_2,42_ = 1.70, p = 0.19), likely due to large interindividual variability (Figure S1E). Nevertheless, scores exhibited a downward trend from the control group (0.53 ± 0.06) to the non-apraxic group (0.47 ± 0.06) and then to the apraxic group (0.37 ± 0.06).

### Spatial working memory was not impaired in individuals with apraxia

Spatial working memory was assessed using the Corsi task in 10 neurotypical participants, 15 participants without apraxia, and 15 participants with apraxia. Similar to the results of the Raven’s Progressive Matrices task, there were no statistically significant differences observed among the three groups (F_2,37_ = 1.72, p = 0.19; Figure S1F), but a gradual decline in performance was noted from the control group (43.60 ± 4.97) to the non-apraxic group (38.87 ± 4.33) and to the apraxic group (31.07 ± 4.02).

### Individuals with apraxia had worse stroke severity

Stroke severity as assessed by the NIH stroke scale score was obtained for all but one participant with left hemisphere stroke (who was in the apraxic group). Stroke severity was significantly worse in the apraxic group than in the non-apraxic group (mean difference = 2.47 ± 1.19, t_24.214_ = 2.07, p = 0.049, 95% CI = [0.01, 4.92], Cohen’s d = 0.76; Figure S1G). This finding highlights the importance of using partial correlations in the sparse network analysis to distinguish true relationships across tasks from apparent relationships driven primarily by overall stroke severity.

**Figure S2:**
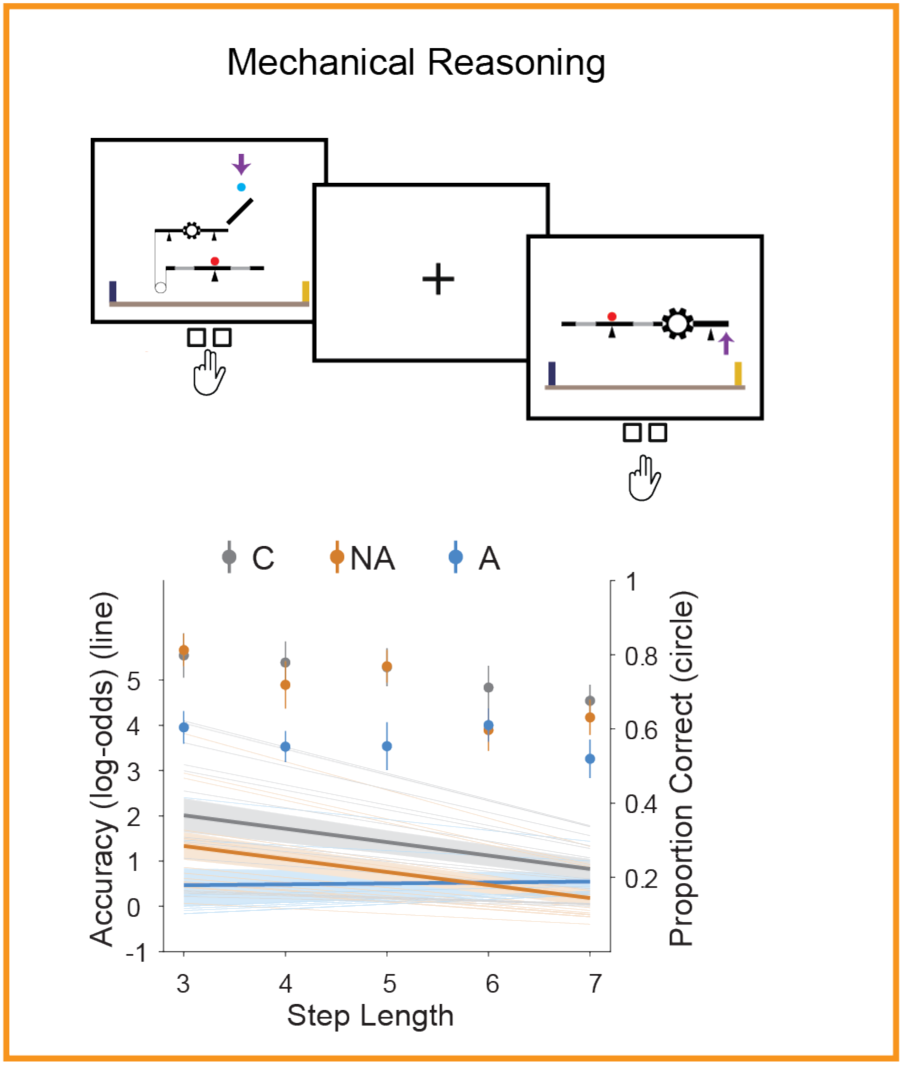
Performance in the Rube Goldberg task. A generalized mixed effect model revealed a significant effect of group on response accuracy (χ_2_^2^ = 6.15, p = 0.046; Figure 1C, Middle), as well as an interaction between group and task difficulty (i.e., step length) (χ_2_^2^ = 9.22, p = 0.01). The main effect of step length was not significant (χ_1_^2^ = 0.05, p = 0.82). Post hoc contrast analyses revealed that response accuracy decreased with difficulty level in the control group (slope = −0.30 ± 0.08, p < 0.0001, 95% CI = [-0.45, - 0.14]) and the non-apraxic group (slope = −0.29 ± 0.07, p < 0.0001, 95% CI = [−0.42, −0.15]]), while it remained stable and near zero in the apraxic group (slope = 0.02 ± 0.09, p = 0.82, 95% CI = [−0.15, −0.19]]). By comparing response accuracy at each step length (Figure 1D), we found that at a step length of 3, the apraxic group was significantly less accurate than the control group (mean difference = −1.55 ± 0.62, p = 0.03, 95% CI = [−3.01, −0.08]. However, there was no significant difference between either the apraxic and non-apraxic groups (mean difference = −0.87 ± 0.60, p = 0.32, 95% CI = [-2.28, 0.55]) or the non-apraxic and control groups (mean difference = −0.68 ± 0.54, p = 0.42, 95% CI = [-1.94, 0.59]). At larger step lengths (i.e., step lengths 4-7), none of the three groups were significantly different from each other (all p > 0.08).

**Figure S3:**
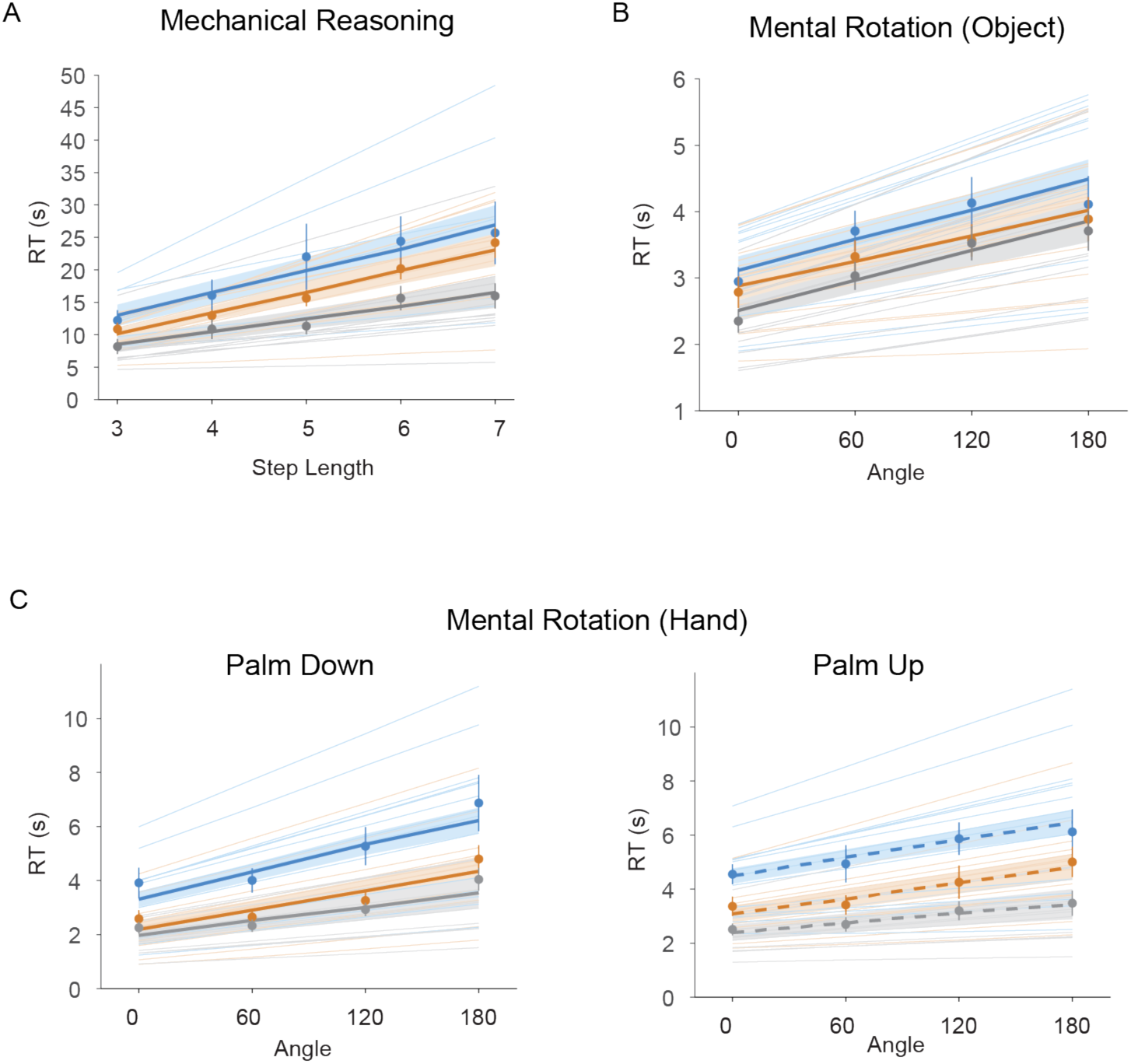
Response time for correct trials across tasks. **A)** In the Rube Goldberg task used to assess mechanical reasoning ability, response time (RT) was affected only by step length. **B)** In the object-based mental rotation task, RT was affected only by rotation angle. **C)** In the mental rotation of hand task used to assess motor imagery, RT was affected by hand image orientation, rotation angle, and group. Specifically, RT was slower for palm-up images compared to palm-down images and increased with larger rotation angles. Importantly, the apraxic group exhibited longer RTs compared to the other two groups.

**Figure S4:**
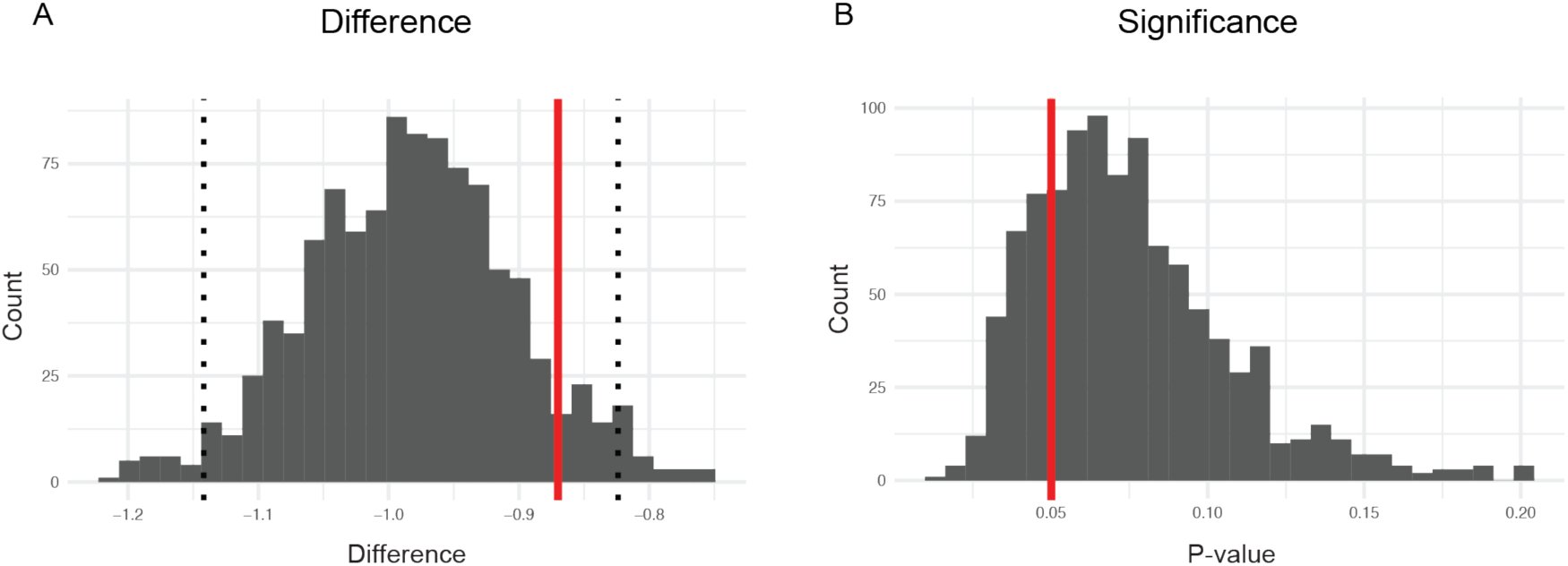
**Differences in mechanical reasoning score between the apraxic and non-apraxic groups when all participants were included**. Our original analysis on mechanical reasoning ability excluded eight out of 16 participants in the apraxic group, two out of 15 participants in the neurotypical control group, and one out of 15 participants in the non-apraxic group, as they failed the criterion test (see Methods). Because they did not proceed to the experimental task, no data were available to measure their mechanical reasoning ability. Here, we assumed that they would randomly guess the response (left or right) if they had have completed the Rube Goldberg task. To simulate their data, we imputed each trial with either a correct response or incorrect response randomly and re-ran the generalized mixed-effect model and post hoc comparisons on the complete dataset from all participants. We repeated this process 1000 times. **A**) The difference between the apraxic group and the non-apraxic group from the post hoc comparisons. Red solid line represents the value from the original analysis and dotted lines represent the significance level at 0.05 determined by the 1000 simulations. The true value within the 95% interval suggests that our original result of no difference between apraxia and non-apraxia was not a special case. **B**) The corresponding p-value of the difference between the apraxic group and the non-apraxic group from the post hoc comparisons. Most simulations yielded non-significant difference between these two groups (i.e., p is not smaller than 0.05). Red solid line represents the significance level at 0.05.

### Zero-order correlations among all measures of interest

For completeness, zero-order correlations among all measures of interest are shown in Figure S5. We note in particular that there were significant correlations between apraxia severity (both pantomime and meaningless imitation) and the scores on novel tool selection and application (all ρ > 0.44; p < 0.05). Strong correlations were also observed between motor imagery and pantomime (ρ = 0.60; p = 0.0059), as well as between motor imagery and meaningless imitation (ρ = 0.56; p = 0.0097). These results are consistent with prior work stressing that deficits in novel tool selection/application (Goldenberg, 2013b) and motor imagery (Buxbaum et al., 2005; Schwoebel et al., 2004; Sirigu et al., 1996) are commonly observed in people with apraxia. We also observed a significant correlation between meaningless imitation and mechanical reasoning (ρ = 0.40; p = 0.0459). Critically, unlike novel tool selection, mechanical reasoning did not show a significant correlation with pantomimed tool-use (ρ = 0.28; p = 0.1542), despite the fact that both novel tool selection and mechanical reasoning were found to be associated with general reasoning and potentially with spatial working memory. Moreover, motor imagery and the mental rotation of objects were not correlated to either novel tool selection or mechanical reasoning.

**Figure S5:**
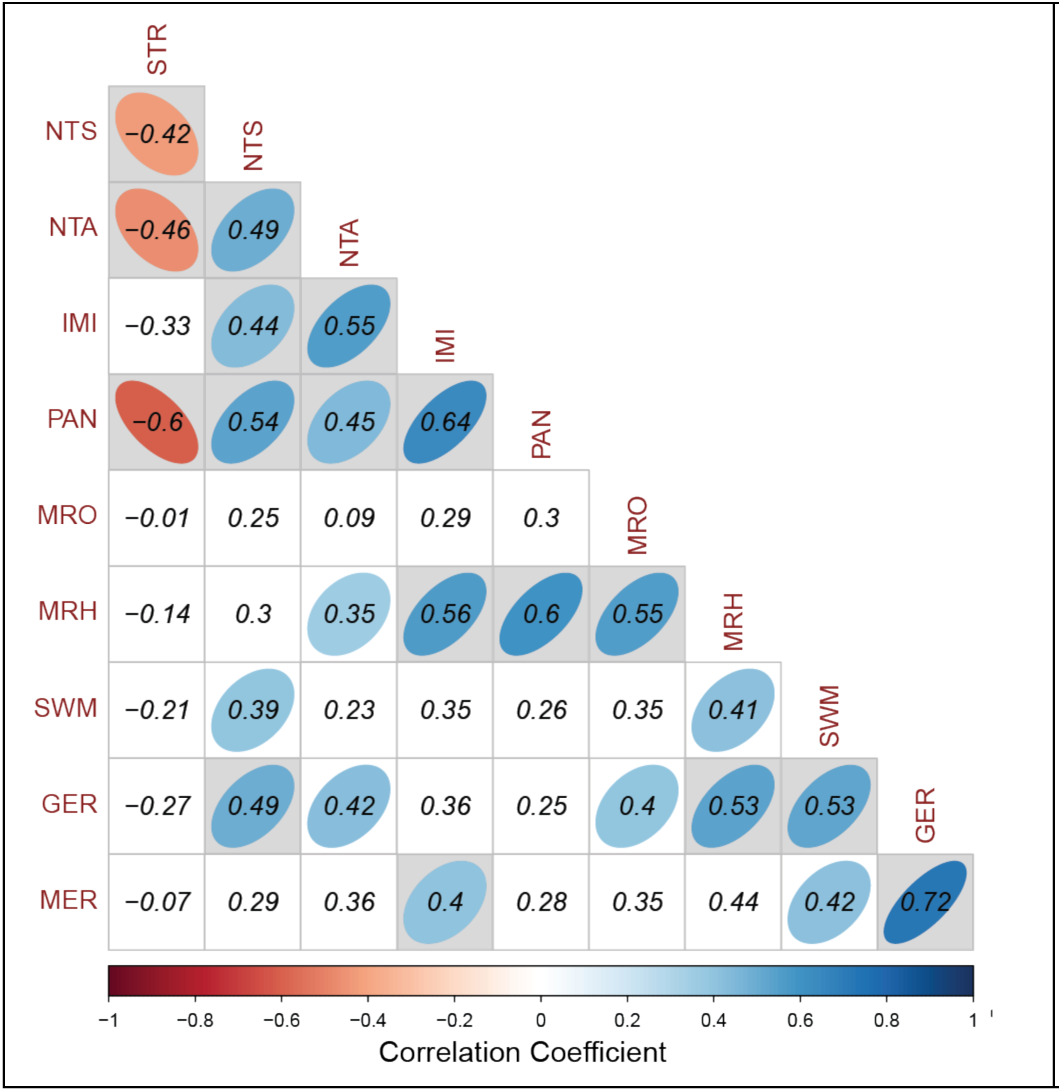
Zero-order correlations among all task scores. **A)** The correlation matrix among the scores of various cognitive tasks including the novel tool and Rube Goldberg tasks. Values are zero-order correlation coefficients between a pair of variables. Cells with an ellipse reflect correlations that were statistically significant (α = 0.05 without multiple comparison correction). Grey cells are correlations that were statistically significant after the False Discovery Rate (FDR) correction. STR: Stroke severity; NTS: Novel tool selection; NTA: Novel tool application; IMI: Meaningless imitation; PAN: Pantomime; MRO: Object-based mental rotation; MRH: mental rotation of hands (motor imagery); SWM: Spatial working memory; GER: General reasoning; MER: Mechanical reasoning.

**Figure S6:**
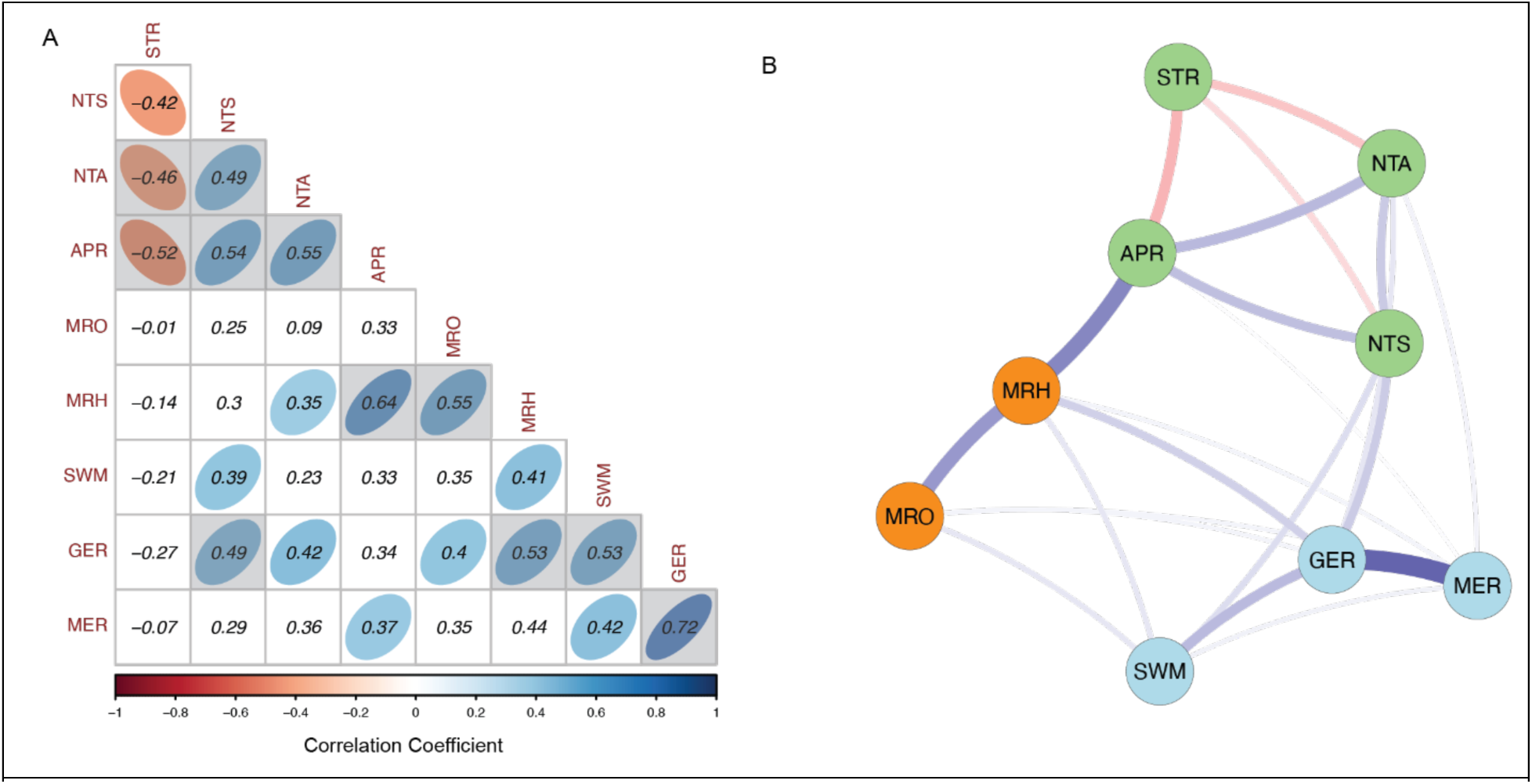
Correlation matrix and Latent structure remained unchanged when combining pantomimed tool-use and meaningless imitation as a single measure of apraxia severity. **A)** The zero-order correlations matrix among the score of various cognitive tasks including the novel tool and Rube Goldberg tasks. Values are zero-order correlation coefficients between a pair of variables. Cells with an ellipse reflects correlations that were statistically significant (α = 0.05 without multiple comparison correction). Grey cells are correlations that were statistically significant after the False Discovery Rate (FDR) correction. **B)** A sparse network among variables established through the graphical lasso analysis. We identified three distinct clusters of tasks (green, orange, and blue circles). Thickness of lines: magnitude of partial correlation. Blue lines: positive partial correlation; Red lines: negative partial correlation. Both the zero-order correlations and the latent structure remained the same as our original analysis with pantomimed tool-use and meaningless imitation as separate measures. APR: apraxia severity; NTS: Novel tool selection; STR: NIH Stroke scale; NTA: Novel tool application; MRO: Mental rotation of object; MRH: Mental rotation of hand; SWM: Spatial working memory; GER: General reasoning; MER: Mechanical Reasoning.

### Detailed description of the Sparse Network (*Figure 2*)

Our network analysis revealed a global structure among all tasks of interest. In addition to the two separate clusters of mechanical reasoning and novel tool selection (see main results), we observed a third cluster reflecting mental rotation and motor imagery, although this community was less stable as it often merged into either of the other two communities. In particular, among all bootstrapped and stimulated networks with more than one cluster, 50% percent of the bootstrapped samples and 73.4% of the simulated samples exhibited only two clusters, where object-based mental rotation was typically merged into the reasoning cluster, while motor imagery (i.e., mental rotation of hands) was equally likely to be grouped with either the reasoning or tool-use/praxis communities (Figure S8B &10B).

Beyond this global organization, the resulting sparse network revealed a subset of crucial tasks and showed rich signals of their direct relationships. Motor imagery, pantomime, and general reasoning, with strong betweenness measures, served as the key nodes that bridged distinct clusters (Figure S7). In particular, motor imagery was strongly connected to pantomime and meaningless imitation, bridging the cluster of tool-use measures and the cluster of mental rotation. This connection was consistently observed in the original network and the bootstrapped and simulated networks (Figures S8-S11). On the other hand, general reasoning was consistently connected to novel tool selection, serving as a bridge connecting the two clusters of tool-use and high-level reasoning functions (Figures S8-S11).

**Figure S7:**
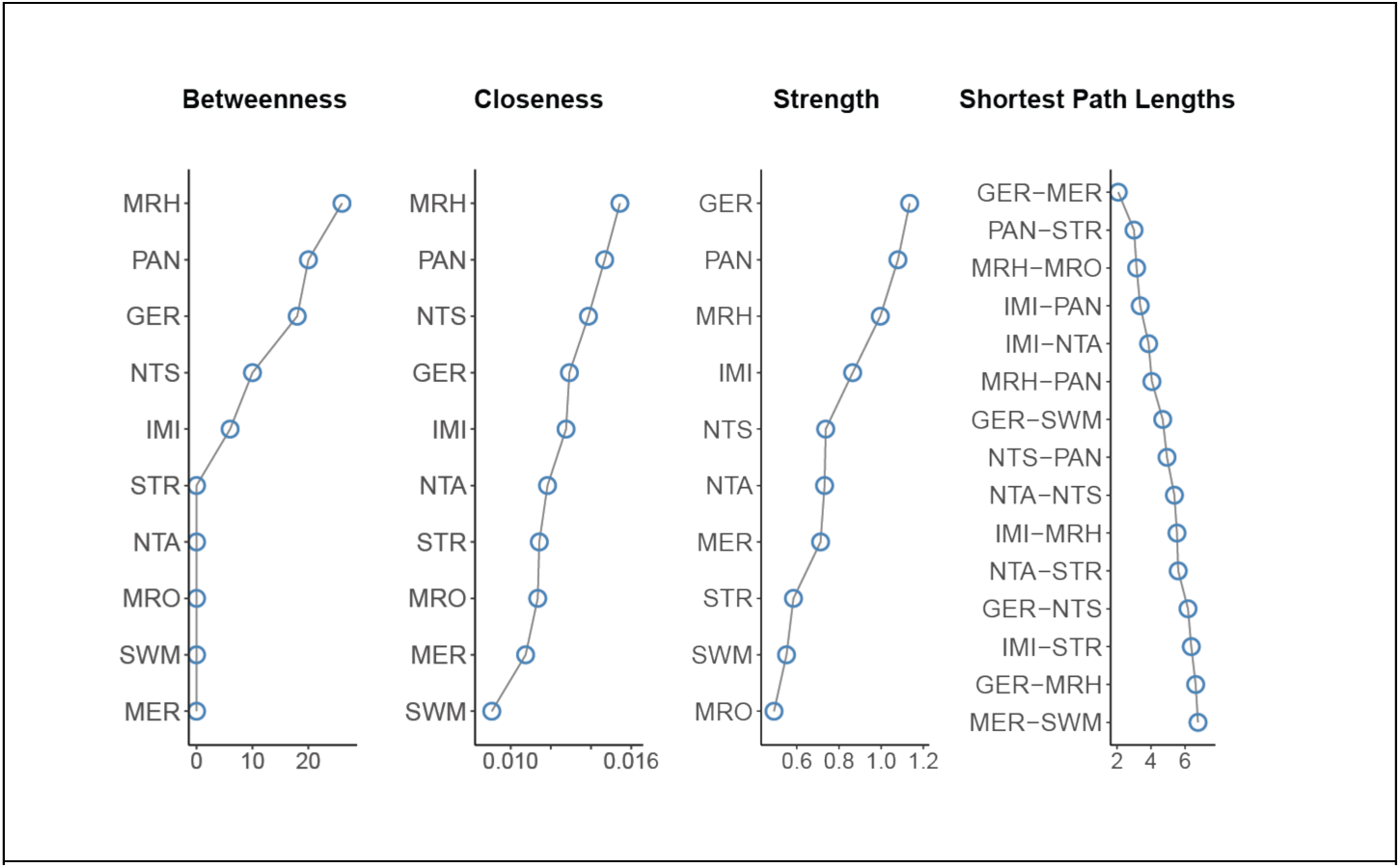
Centrality measures of the original network. The importance of nodes in the network was revealed by centrality measures including Betweenness, closeness, and strength. Note these metrics were used for exploratory comparison, not for formal statistical inference. Only the top 15 pairs of tasks are shown. Higher betweenness score corresponds to more important roles as bridges between communities in the network. Closeness centrality reflects how globally connected a task is to all other tasks in the network, based on the shortest paths. Strength centrality measures the total magnitude of direct connections a task has with all other tasks. The shortest path length between a pair of nodes highlights the direct relationship between nodes. Lower scores suggest a more direct relationship (e.g., faster travel from one task to the other). Note these measurements are for descriptive comparisons, not for statistical inferences.

**Figure S8:**
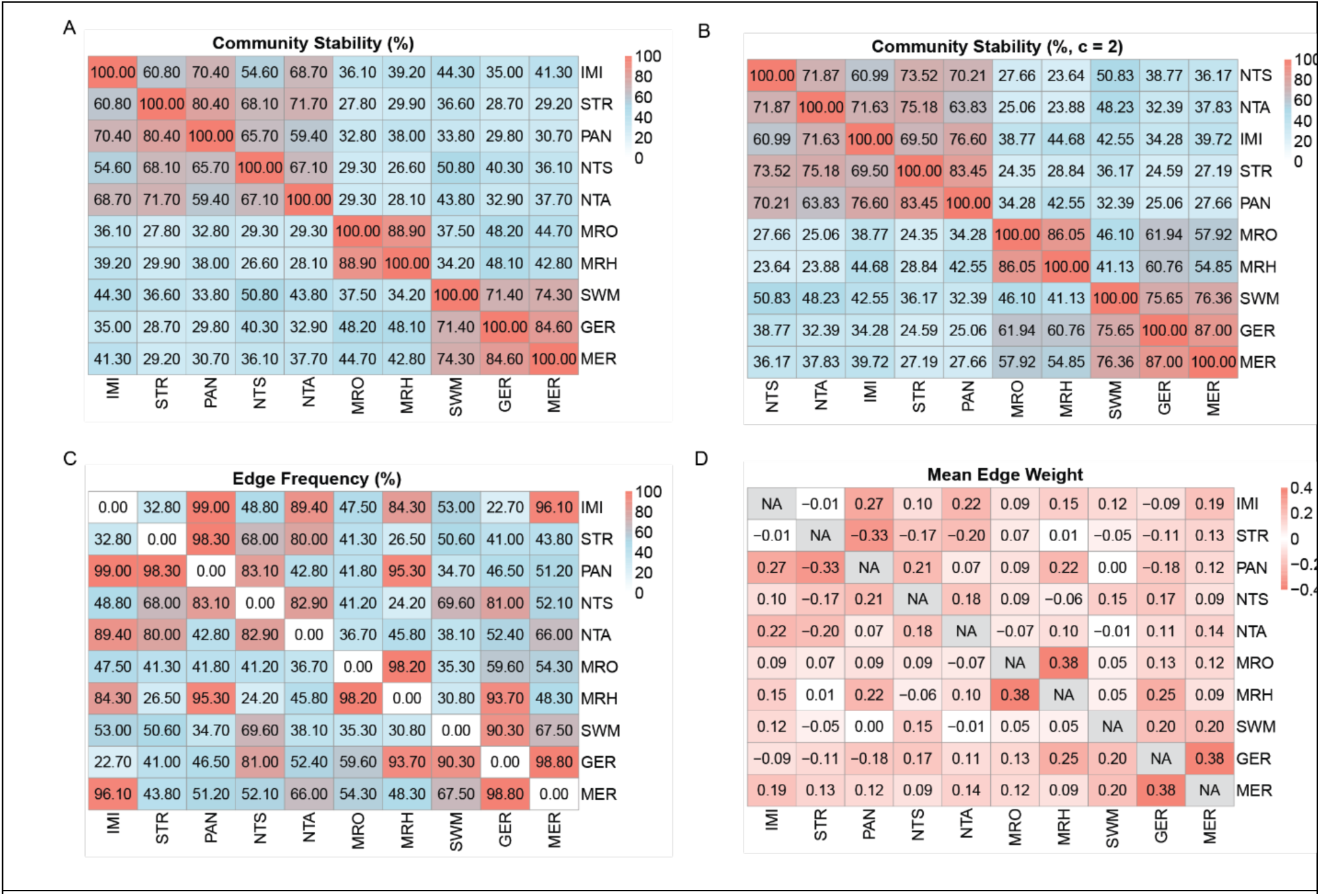
Network stability across bootstrap samples (n = 1000). **A**) Percentage of networks in which each pair of tasks was clustered into the same community. **B**) Percentage of networks in which each task pair was clustered into the same community when there were only two clusters. **C**) Percentage of networks in which connections between task pairs survived after graphic lasso regularization. NTS showed reliable connections to PAN, NTA, and GER (> 80%), while MER exhibited reliable connections to GER and IMI. **D**) Mean connection weight (if survived) between each pair of tasks.

**Figure S10:**
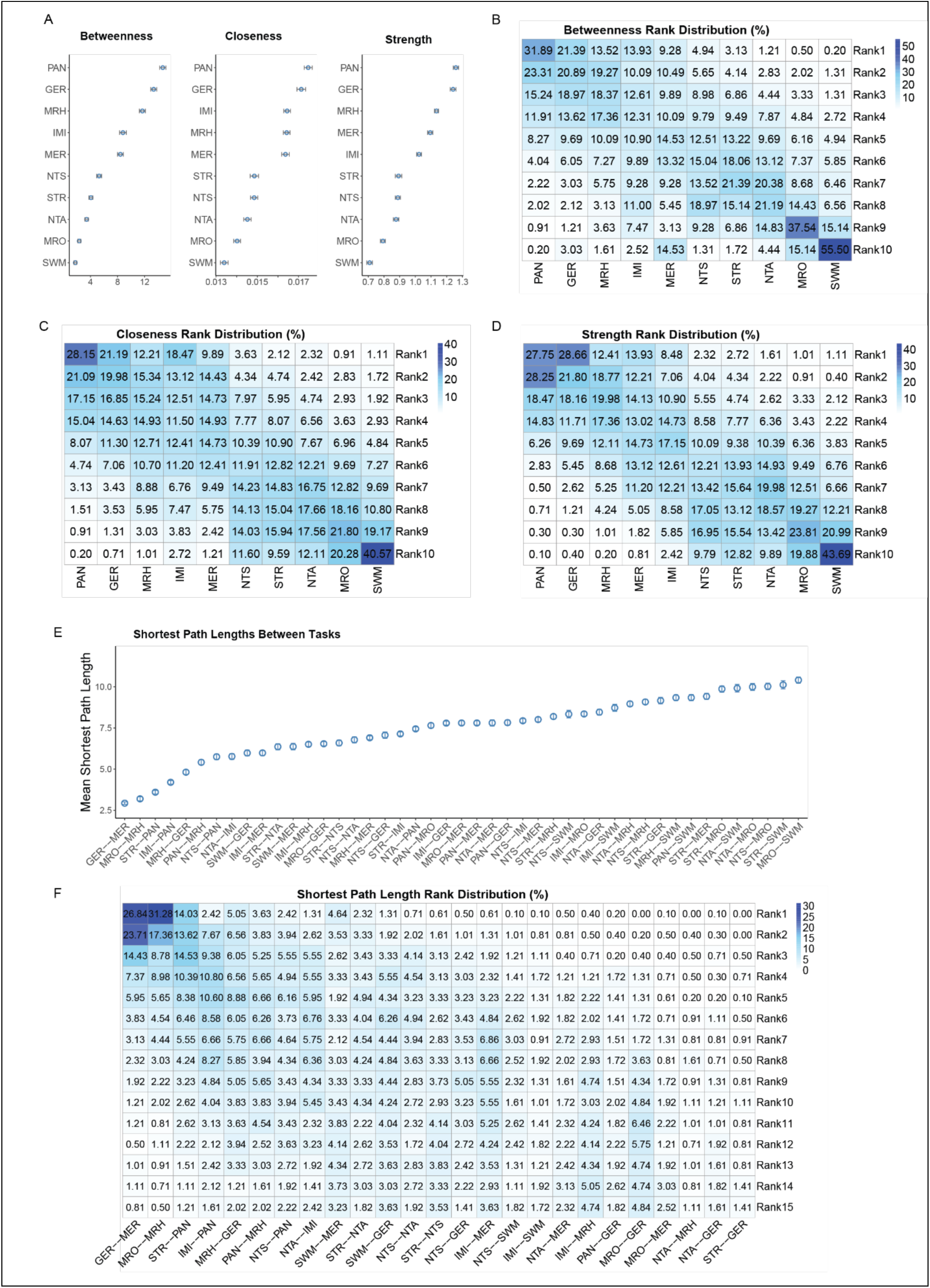
Network stability across simulated samples (n = 1000). **A**) Percentage of networks in which each pair of tasks was assigned to the same community. **B**) Percentage of networks in which each task pair was clustered together under a two-community solution. **C**) Percentage of networks in which the connection between task pairs persisted after graphical lasso regularization. **D**) Mean connection weight (if survived) between each pair of tasks. The edge frequencies and edge weights were similar to those in the bootstrap samples. A notable exception was the connection between NTS and IMI, which appeared consistently but had a low mean weight, suggesting a potentially non-meaningful relationship.

**Figure S9:**
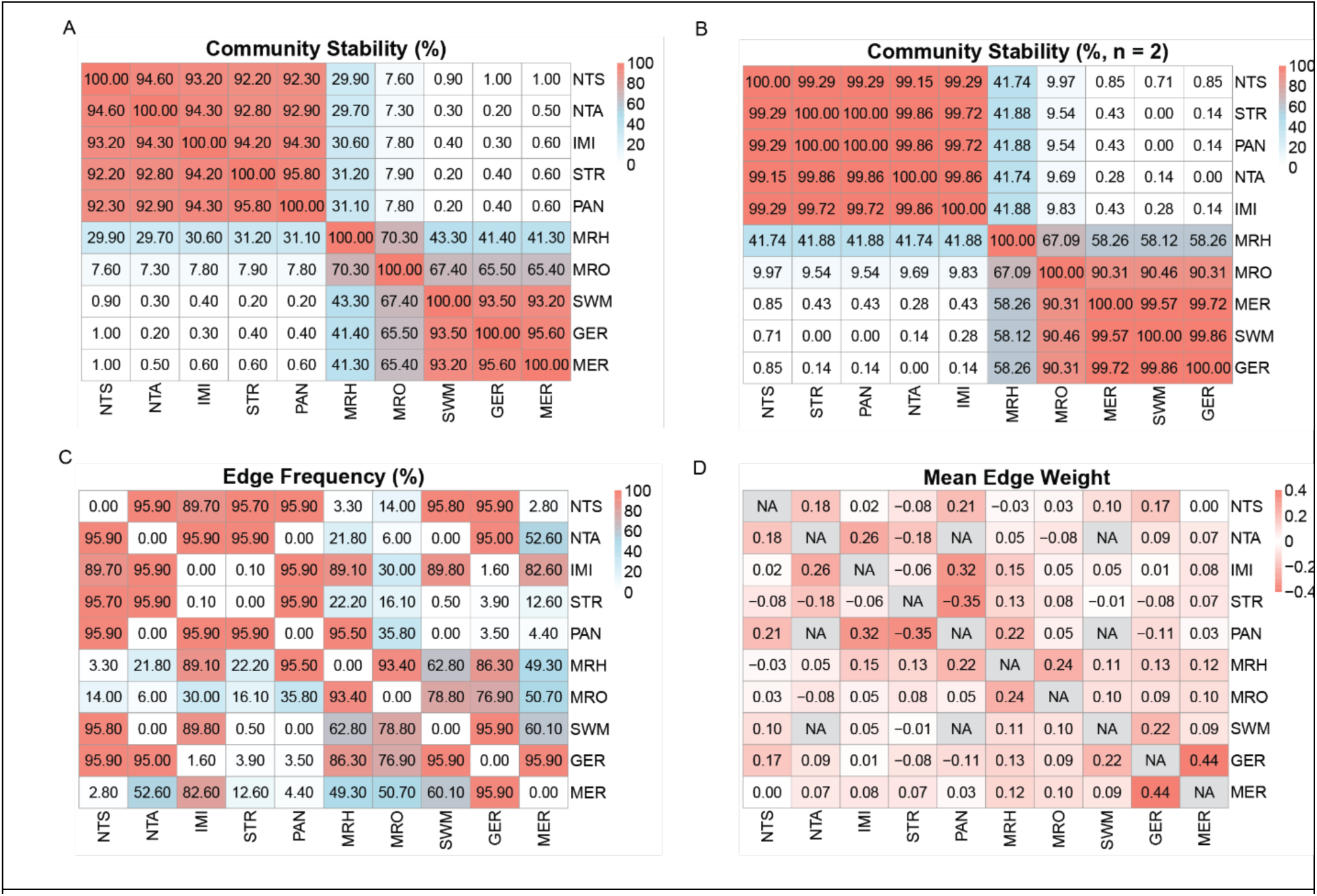
Centrality and ranking measure across bootstrap samples (n =1000). **A**) Betweenness, closeness, strength centrality values of each task. Error bars represent 95% confidence intervals. **B**) Ranking matrix of betweenness, **C**) Closeness, and **D**) Strength. Each cell reflects the percentage of bootstrap networks in which a task held a specific rank (e.g., rank 1 reflects highest centrality). Task names were ordered by the cumulative percentage of being in the top 4 ranks for each measure. E) Mean shortest path length across all sampled networks. **F**) Ranking matrix of the shortest path lengths. Only the top 15 pairs (out of 45 pairs) are shown.

**Figure S11:**
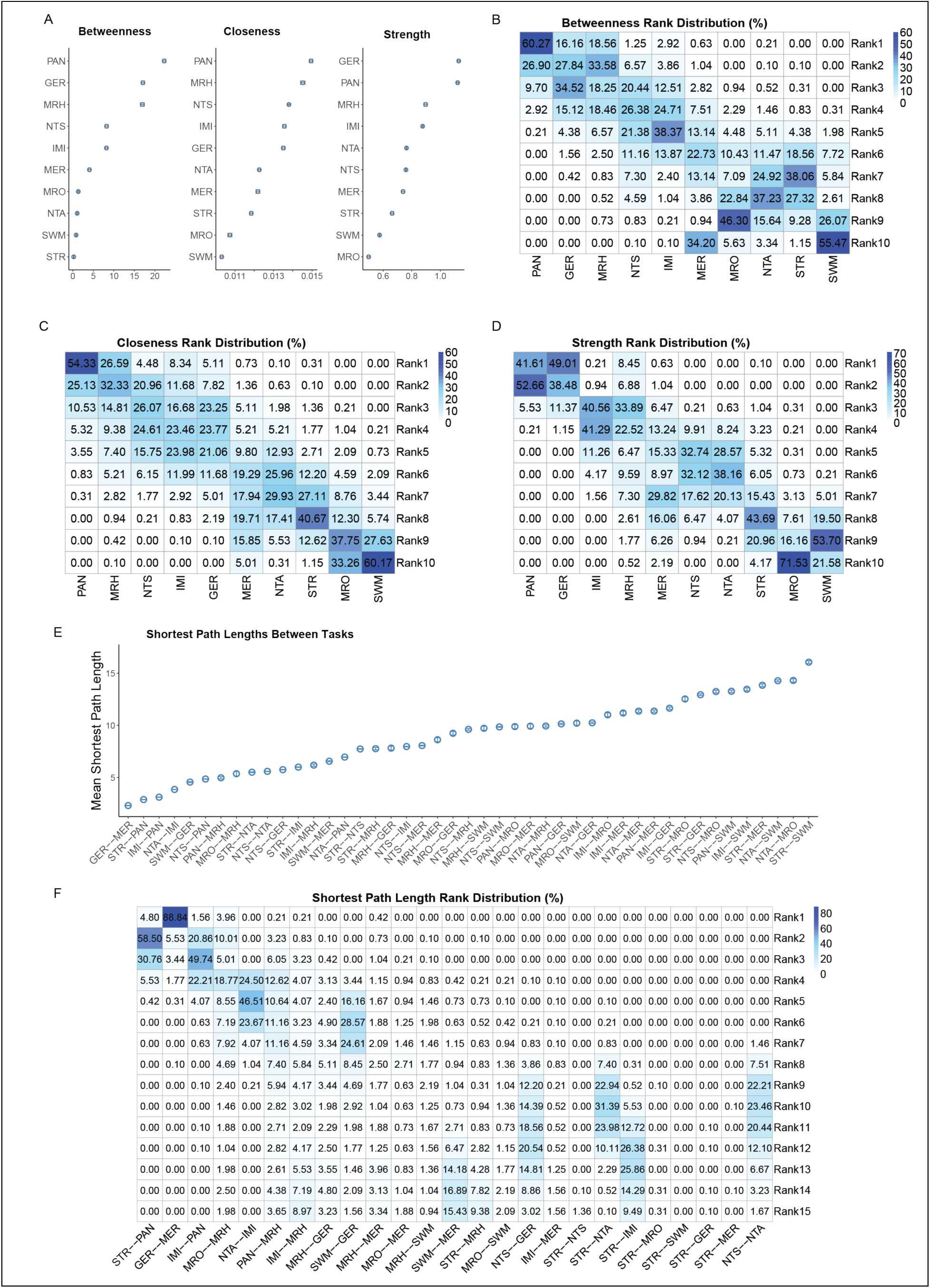
Centrality and ranking measure across simulated samples (n =1000). **A**) Betweenness, closeness, strength centrality values of each task. Error bars represent 95% confidence intervals. **B**) Ranking matrix of betweenness, **C**) Closeness, and **D**) Strength. Each cell indicates the percentage of simulated networks in which a task held a specific rank. Task names were ordered by the cumulative percentage of being at top 4 ranks for each measure. E) Mean shortest path length between tasks across all sampled networks. **F**) Ranking matrix of the shortest path lengths. Only the top 15 pairs out of 45 pairs are shown.

**Figure S12:**
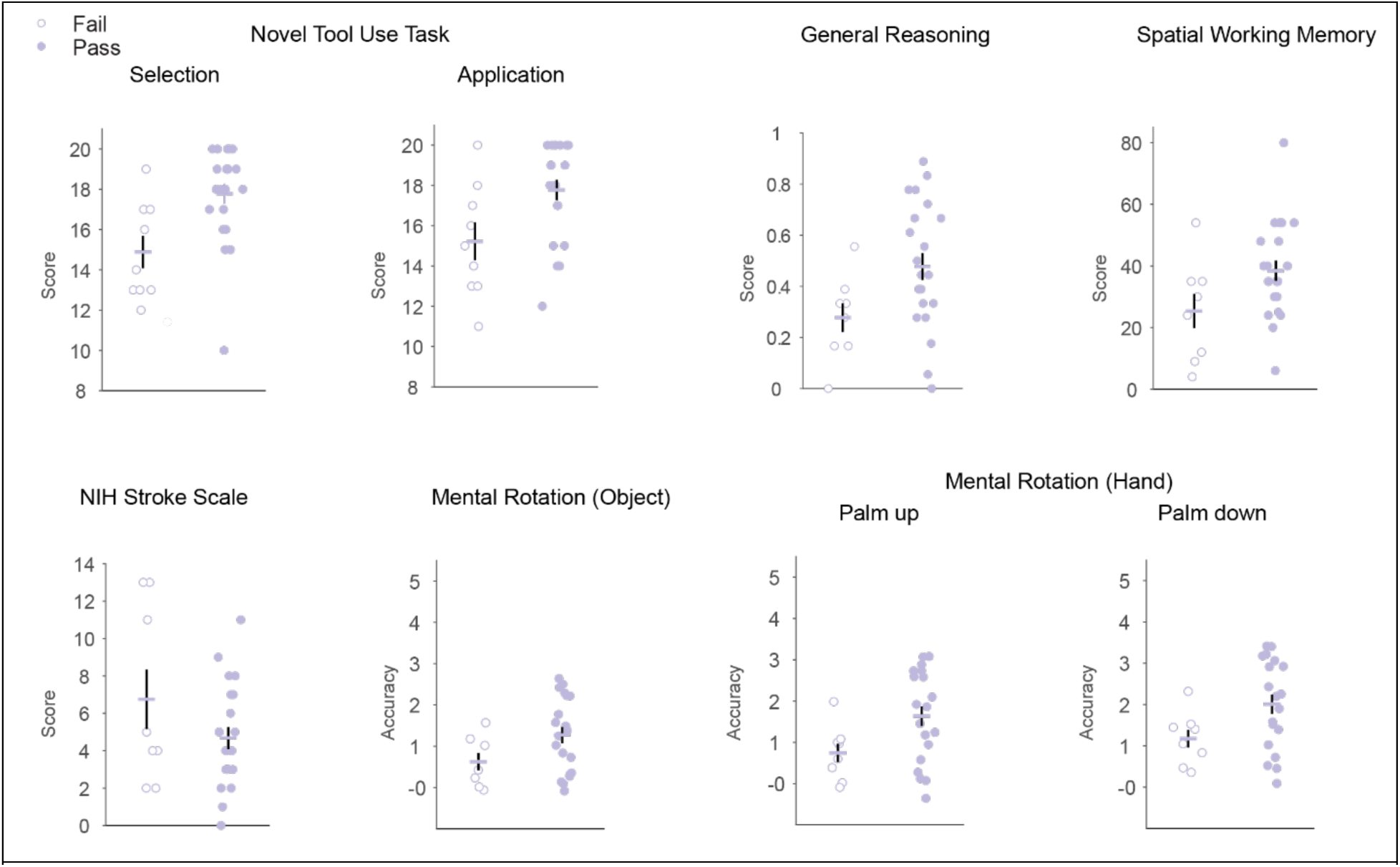
Performance of individuals who failed and who passed the criterion test in the Rube Goldberg Task. Among all participants with left hemisphere stroke, individuals who failed the criterion task exhibited performance comparable to the lower-performing half of those who passed the criterion task, across all tasks except overall stroke severity.

## Notes

### Competing Interest Statement

The authors have declared no competing interest.

